# Function aligns with geometry in locally connected neuronal networks

**DOI:** 10.1101/2025.08.08.669348

**Authors:** Antoine Légaré, Olivier Ribordy, Paul De Koninck, Antoine Allard, Patrick Desrosiers

## Abstract

The geometry of the brain imposes fundamental constraints on neuronal network organization and dynamics, yet how these constraints give rise to observed patterns of brain activity remains unclear. Here, we investigate how geometric eigenmodes relate to functional connectivity gradients in three-dimensional neural systems using a combination of generative network simulations and cellular-resolution calcium imaging in larval zebrafish. We show that functional connectivity gradients emerging from network activity naturally align with the geometric eigenmodes of the underlying spatial embedding when connectivity is predominantly local. By systematically increasing the prevalence of long-range connections, we reveal a robust geometry-function correspondence that progressively deteriorates as local connectivity is disrupted. We then show that spatial filtering can artificially imprint geometric patterns on functional gradients, highlighting an important methodological confound. To validate our computational results, we conduct volumetric calcium imaging experiments at cellular resolution in the optic tectum of zebrafish larvae, uncovering functional gradients that closely align with geometric eigenmodes. As predicted from simulations, the eigenmode-gradient mapping exhibits a cutoff point that quantitatively reflects the spatial extent of the region’s connectivity kernel, inferred from single-neuron morphologies. This geometry-functional alignment disappears at brain-wide scale, where long-range connections are more prevalent. Our findings demonstrate how short-range anatomical connectivity anchors functional connectivity gradients to the brain’s geometry.

## Introduction

Understanding what guides the flow of brain-wide activity remains a central challenge in systems neuroscience. At any moment, several factors influence brain dynamics, including external sensory inputs and ongoing internal demands^1^. Yet beneath these transient influences lies a more fundamental constraint: the brain’s intrinsic connectivity^2^. As in many natural systems, structure and function in the brain are deeply intertwined. Anatomical wiring diagrams predict coordinated activity across neurons^3;4;5^ or brain regions^6^, while the three-dimensional arrangement of neural populations across gyri, sulci, and subcortical nuclei mirrors their functional specialization^7^. A comprehensive account of how activity flows through the brain must therefore integrate both its network architecture and its spatial embedding—its geometry.

In human neuroimaging, network science has been widely employed to model both anatomical connectivity and brain dynamics^8^. One of the most common network-based descriptions of brain activity is functional connectivity (FC), typically defined as the correlation between activity time series from distinct neurons or brain regions. FC provides a compact statistical summary of interactions between neuronal populations over extended timescales, but it does not describe rapid, moment-to-moment fluctuations of the brain’s activity^9^. An influential extension of FC analysis is the extraction of functional connectivity gradients — eigenvectors derived from the FC matrix that capture the dominant axes of variation in connectivity^10;11;12^. Gradients have become useful tools as they coincide with the spatial organization of brain functions, not only in humans but also across species, including mice and zebrafish^6;13^.

Although connectivity gradients inherit the temporally static nature of the FC matrix, the resulting spatial configurations can provide insights into underlying dynamical processes. Recent work has shown that FC is driven by propagating waves of hemodynamic activity, which diffuse across canonical resting-state networks organized along the functional gradients^14;15^. Consistent with macroscale observations, waves of neuronal activity have been observed across cortical and subcortical systems in a wide range of species and recording modalities^15;16;17;18^. While their mechanistic origins and computational roles remain debated, such waves are thought to arise naturally from fundamental architectural features of brain networks, including conduction delays and the predominance of short-range connectivity^19^.

A hallmark property of brain networks is wiring-length minimization, which favors dense local connectivity^20^. Although long-range projections are essential to coordinate global dynamics^21;22;23^, their relative sparsity suggests that local interactions dominate the generation of neural activity. Neural Field Theory (NFT) formalizes this principle by modeling brain activity as the propagation of electrical waves across a continuous neural sheet^24;25;26^ governed by distance-dependent connectivity, often approximated by an exponential decay rule (EDR)^27^. Building on this framework, Pang and colleagues recently tested the hypothesis that geometry itself constrains brain dynamics beyond the effects of anatomical connectivity^28^. Using MRI, they demonstrated that eigenmodes of the Laplace–Beltrami operator (LBO) applied to cortical geometry provide a parsimonious basis to reconstruct brain activation patterns. Strikingly, they further reported a near one-to-one correspondence between geometric eigenmodes and FC gradients in subcortical nuclei, suggesting that shape alone can predict the dominant axes of functional organization^28^. These findings highlight a potentially fundamental role for geometry in shaping brain-wide functional organization, yet the generative mechanisms linking geometry, connectivity, and activity remain incompletely understood. Moreover, the limited spatial resolution of fMRI—combined with common preprocessing steps such as spatial filtering—can bias gradient analyses^29^, underscoring the need for validation in alternative experimental systems where activity is measured at cellular resolution and without such confounds.

In this study, we combine numerical simulations and optical imaging at cellular resolution to investigate the link between FC gradients and geometric eigenmodes in brain networks. Using dynamical models of neuronal networks embedded in various 3D shapes, we show that the correspondence between FC gradients and geometric eigenmodes emerges naturally from short-range connectivity and local correlations, independent of specific spatiotemporal dynamics. We further demonstrate that this correspondence deteriorates as local connectivity is disrupted, revealing a wavelength-dependent breakdown of the eigenmode-gradient correlations that serves as a sensitive readout of the underlying network’s connectivity kernel. To validate our numerical findings and extend previous neuroimaging results^28^, we perform two-photon calcium imaging in the larval zebrafish optic tectum, a brain structure with local recurrent connectivity. We observe functional gradients that coincide with the geometric eigenmodes of the tectum and, consistent with our simulations, the cutoff point of this mapping predicts the decay parameter of the exponential law that governs tectal connectivity, which we estimated from reconstructed neuronal morphologies. Altogether, our results identify a simple generative mechanism for the emergence of geometric connectivity gradients and statistical patterns in the brain, underscoring the prevailing influence of geometry on brain wiring and functional organization.

## Results

Throughout this study, we compare geometric eigenmodes and FC gradients (hereafter referred to as eigenmodes and gradients, respectively) in neuronal populations embedded within three-dimensional volumes of various shapes. Geometric eigenmodes are the eigenvectors of the Laplace–Beltrami operator (LBO, Methods), and capture spatial patterns, analogous to vibrational modes, that arise from the shape of the embedding. FC gradients are computed from pairwise correlation between nodes activity using diffusion maps^30^ and represent principal axes of variation in network connectivity (Methods). When applied to human brain connectivity data, FC gradients spatially map onto different brain functions, providing a data-driven measure of functional organization^10^. Importantly, eigenmodes and gradients constitute two independent characterizations of the same system: the former are determined exclusively by geometry, while the latter are derived from neuronal activity. Their correspondence therefore quantifies the extent to which functional organization aligns with the network’s embedding geometry.

### Eigenmodes and gradients in locally connected networks

To explore the relationship between gradients and eigenmodes, we first simulated neuronal population dynamics in various three-dimensional geometries, understood here as Riemannian manifolds embedded in Euclidean space. Neurons were assigned uniformly random coordinates within each geometry, then connected according to an exponential decay rule *(EDR)*^27^, that is, with a probability 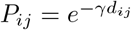 that decays with distance *d*_*ij*_ between neurons *i* and *j*. The parameter *γ* > 0 controls the decay rate, with high values yielding locally connected networks, and low values increasing the likelihood of long-range connections. We will also refer to the inverse of this parameter, 1*/γ*, as the characteristic length of the network’s connections. As a first case study, we generated a simple 3D ellipsoid with *N* = 2500 locally connected neurons and *γ* = 25, yielding connections that on average spanned roughly 10% of the ellipsoid’s length (Fig. 1**a**). We randomly assigned a weight to each connection then simulated simple chaotic firing rate dynamics^31^ (model 1) resulting from nonlinear excitatory and inhibitory interactions between connected neurons (Fig. 1**b**, Methods). As the network density varied with the random sampling process and the *γ* parameter, we also normalized synaptic weights by the network’s density *ρ*, ensuring consistent weight distributions over subsequent simulations. We then computed an average correlation matrix *C* (representing functional connectivity, Fig.1 **c**) from the firing rates of neurons by averaging the absolute value of the pairwise correlations from 50 simulations, throughout which node coordinates and connections were preserved, but weights and initial conditions were drawn randomly each time. Firing rate correlations decayed monotonously and exponentially with distance (Fig.1 **d**), mirroring the underlying wiring rule and indicating spatially correlated activity within the geometry.

**Figure 1.**
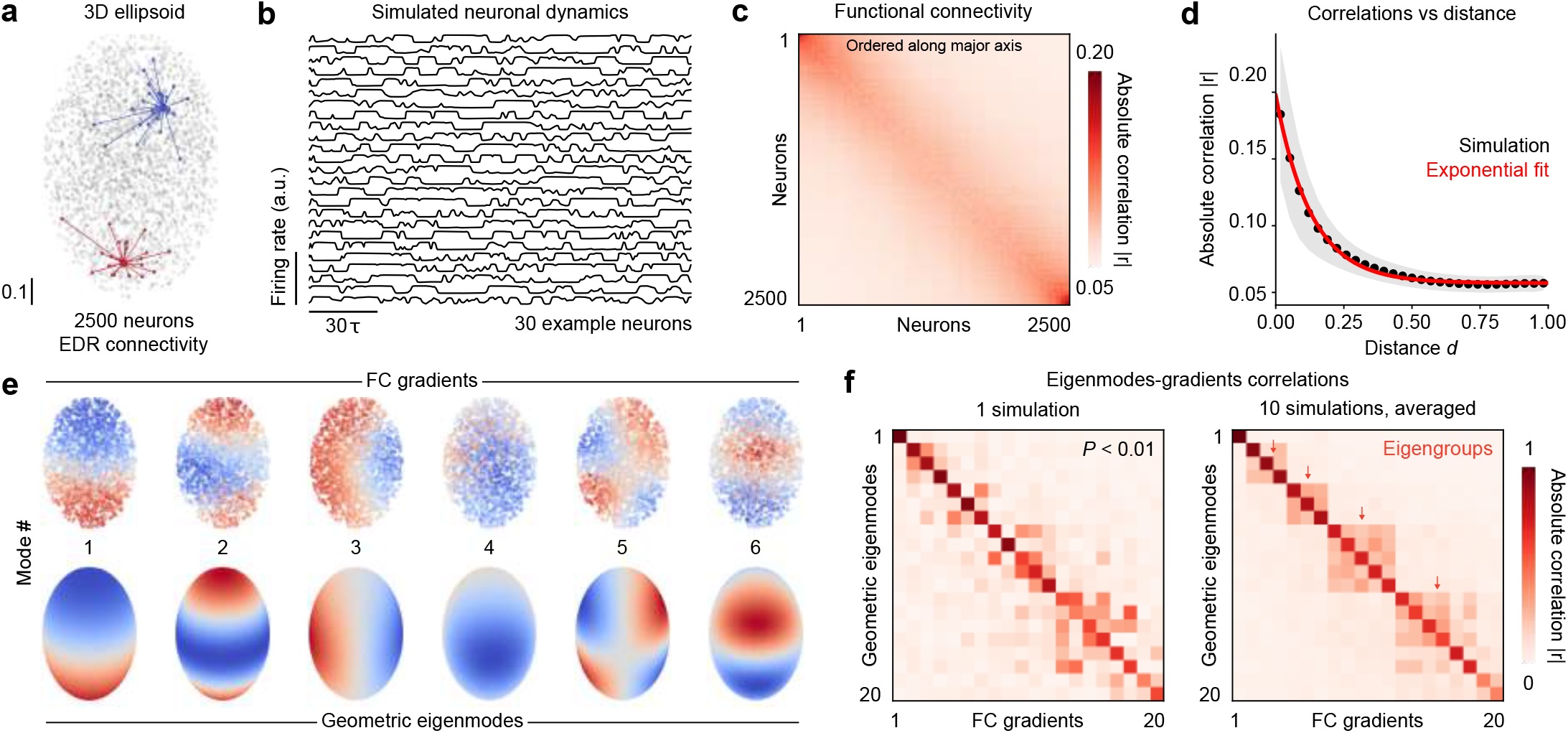
Geometric gradients emerge in firing rate networks with local connectivity. (**a**) Centroids of 2500 neurons in a 3D ellipsoid; example local connectivity profiles with *γ* = 25 are highlighted in red and blue. (**b**) Firing rate dynamics of a random subset of 50 neurons from one example simulation; *τ* indicates the time constant of firing rate neurons. (**c**) Average correlation matrix from 50 simulations; neurons are ordered along the major (vertical) axis of the ellipsoid. (**d**) Relationship between correlation and distance from one representative simulation, taken from the previous panel. The black dots indicate the average per distance bin, and the shaded region indicates standard deviation. A red exponential fit is overlayed for comparison. (**e**) Top row, six functional gradients obtained using diffusion eigenmap, ordered based on their correlation with geometric modes; bottom row, first six geometric eigenmodes of the ellipsoid, with the first constant eigenmode excluded. (**f**) Left, absolute correlation matrix of the first 20 eigenmode-gradient pairs, after functional gradients have been optimally mapped to geometric eigenmodes; right, average of 10 mode correlation matrices, revealing a diagonal block structure corresponding to eigengroups (indicated by arrows).

Next, we computed FC gradients from the averaged *C* matrix using the diffusion maps method^30^, revealing smooth spatial functions when projecting the eigenvectors onto network nodes (Fig.1 **e**, top row, Methods). Independently, we numerically evaluated the geometric eigenmodes of the ellipsoid (Fig.1 **e**, bottom row, Methods), and correlated them with functional gradients after finding an optimal mapping between both mode ensembles (Fig.1 **f** left, 20 mode pairs, Methods). We observed a strong correspondence between eigenmodes and gradients after mapping, with correlations often exceeding |*r*| = 0.9 (average |*r*| = 0.737, Fig.1 **f**). These correlations were also specific, as off-diagonal values were substantially lower (average |*r*| = 0.069), indicating very little spurious correlations between unpaired modes. This correspondence was statistically significant, as spatially permuted eigenmodes yielded lower diagonal correlations and higher off-diagonal correlations with gradients (*P* < 0.01, 100 spatial surrogate sets with preserved autocorrelation^32^, Methods). Averaging eigenmode-gradient correlations over 10 different random coordinate sets and simulated EDR network dynamics further revealed off-diagonal blocks corresponding to rotationally symmetric eigenmodes (eigengroups, Fig.1 **f**, right). These within-group correlations arise from arbitrary rotational misalignments between gradients and eigenmodes induced by random coordinate sampling. Beyond these groups, correlations between unpaired eigenmodes and gradients were close to zero.

Altogether, this first analysis demonstrates that *geometric gradients* —defined here as FC gradients that correlate strongly and specifically with geometric eigenmodes—emerge readily from the activity of locally connected networks. This conclusion does not depend on the choice of geometry (see Supp. Fig. S1 for more complex geometries), nor does it depend on the choice of gradient derivation methodology (see Supp. Fig. S3 for other parameterizations and definitions).

### Geometric gradients emerge from various network dynamics

To further generalize the previous observations, we conducted simulations of EDR networks within the same ellipsoid, but using different dynamical models (Supp. Fig.S2, Methods): model 1 with Dale’s law being enforced on the connectivity matrix^33^ (model 2), Kuramoto-Sakaguchi (KS) oscillators^34^ (model 3), binary stochastic neurons (model 4), and spiking leaky integrate-and-fire (LIF) neurons with simple calcium dynamics^35^ (model 5). Despite very different activity profiles (Supp. Fig.S2 **a**, Supp. Video 1), each model yielded local and spatially decaying correlations (Supp. Fig.S2 **b-c**). Consequently, the gradients calculated from their averaged correlation matrices correlated with the ellipsoid’s geometric eigenmodes (Supp. Fig.S2 **d**). While all dynamics yielded gradients that correlated positively with eigenmodes, some aligned better with geometry. Notably, gradients derived from models 1, 2 and 3 correlated more significantly with geometric eigenmodes than models 4 and 5 (Supp. Fig.S2 **e**). Binary and spiking neurons yielded the lowest correlations with eigenmodes, possibly due to the presence of more spurious long-range correlations or globalized dynamics. Indeed, across all five dynamical models, we observed a strong positive relationship between the strength of the correlation-distance relationship, quantified using Spearman’s ranked correlation, and the quality of the eigenmode-gradient correspondence, quantified by averaging the absolute correlation |*r*| across the first 50 mode pairs (Supp Fig.S2 **f**, Pearson *r* = 0.793). This suggests that geometric gradients are degraded when activity correlations are not as tightly bound to Euclidean distance. Interestingly, the dynamical models used here exhibit spatially disorganized dynamics, rather than coherent, wave-like activity (Supp. Video 1), which has been proposed as a mechanism underlying geometric alignment in functional imaging measurements^28^. These results suggest that wave-like dynamics are not a prerequisite for the emergence of geometric gradients. Rather, the predominant locality of connections and correlations between neurons, regardless of their dynamical regime, is sufficient to drive this phenomenon, as we show in the next two sections.

### Geometric gradients vanish rapidly as connection length increases

We next evaluated how correlations between eigenmodes and gradients depend on network connectivity parameters. We simulated dynamics on networks of *N* = 2500 neurons generated using the EDR wiring rule, while systematically varying the characteristic length 1*/γ*, which controls the probability of long-range connections within the geometry (Fig.2 **a**). We quantified the mean correlation between the first 50 eigenmodes and their corresponding gradients, revealing a peak near *γ*^−1^ ≈ 0.0364 (Fig.2 **b**). At very small 1*/γ* values, corresponding to highly localized connectivity, eigenmode-gradient correlations were reduced. In this regime, however, networks were extremely sparse and failed to reliably sustain rich dynamics, which accounts for the observed decrease in alignment (Fig. 2**b**, left of the peak; Supplementary Fig. S4**a**,**b**). Beyond this sparse regime, increasing 1*/γ* to add more long-range connections quickly reached an optimum, and further increases resulted in a gradual reduction of eigenmode-gradient correlations (Fig. 2**b**, right of the peak). Hence, for finite-sized networks, there exists an optimal regime of predominantly short-range connectivity in which neural dynamics are rich and localized, and geometric gradients emerge. Note that although increasing the characteristic length 1*/γ* increases overall network density, dynamics exhibited stable properties at higher densities (1*/γ* > 0.0333), ruling out the possibility that the loss of geometric alignment arises from qualitative change in network activity (Supplementary Fig. S4**b-f**).

**Figure 2.**
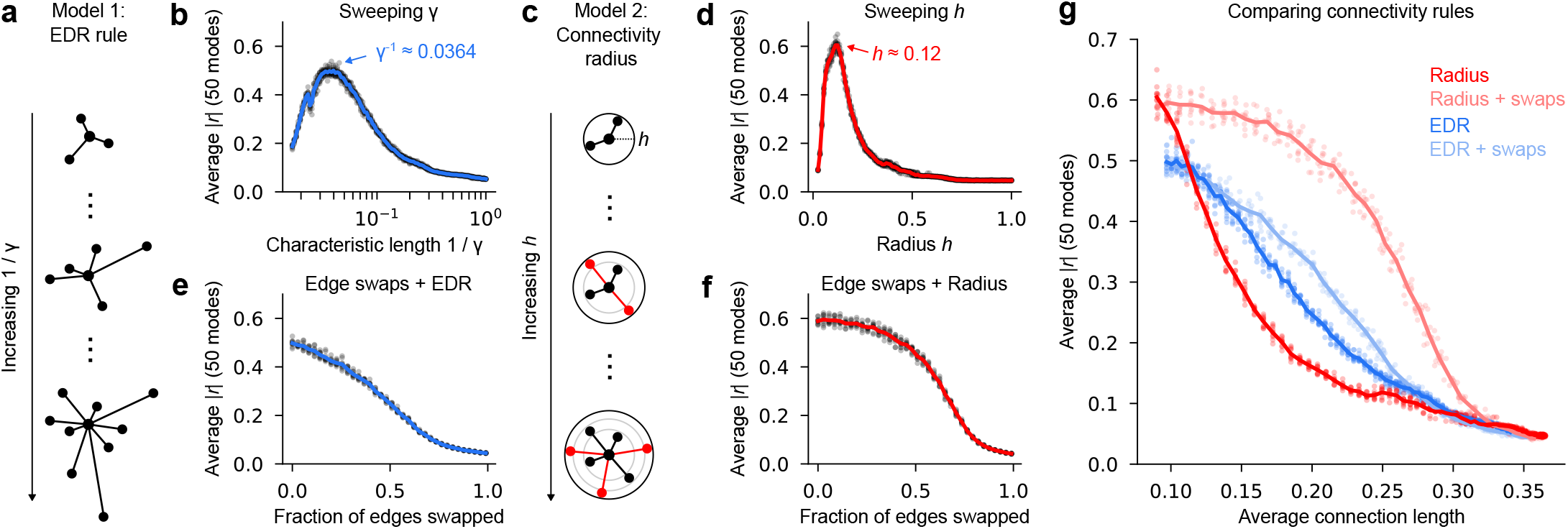
Eigenmode-gradient correlations decrease as the connectivity radius increases. (**a**) Schematization of the EDR model; increasing the characteristic length 1*/γ* increases the probability of long-range connections. (**b**) Absolute eigenmode-gradient correlations |*r*|, averaged over 50 mode pairs for different 1*/γ* values. (**c**) Schematization of the connectivity radius model; increasing the radius *h* deterministically adds new connections, indicated in red. (**d**) Absolute eigenmode-gradient correlations |*r*|, averaged over 50 mode pairs for different *h* values. (**e**) Absolute eigenmode-gradient correlations |*r*|, averaged over 50 mode pairs for different *ρ*_swaps_ values, beginning from EDR networks with 1*/γ* = 0.0364. (**f**) Absolute eigenmode-gradient correlations |*r*|, averaged over 50 mode pairs for different *ρ*_swaps_ values, beginning from radius networks with *h* = 0.125. (**g**) Comparison of the decrease in |*r*| as a function of the network’s average connection length for the four previous models. For all plots, solid lines represent the average, while dots represent individual simulations; 10 simulations per parameter increment.

Because the EDR model probabilistically generates both short- and long-range connections, we next evaluated an alternative wiring rule to isolate the specific contribution of short-range connectivity. We considered a simplified random geometric network model with a hard connectivity threshold. In this model, all neurons separated by a distance *d*_*ij*_ smaller than a fixed radius *h* are connected (Fig. 2**c**). For sufficiently small values of *h* relative to the overall system size, networks are therefore strictly locally connected and devoid of long-range connections. Consistent with the previous results, eigenmode-gradient correlations peaked at small connectivity radii (*h* ≈ 0.12) and decayed abruptly as the radius increased Fig. 2**d**. Peak correlations exceeded those observed under the EDR rule (|*r*| = 0.605 versus |*r*| = 0.500), indicating that the absence of long-range connections in the radius model further enhances the alignment with geometry. In principle, a perfect alignment is expected in the limit *N* → ∞ with vanishing connectivity radius (*h* → 0), as predicted by previous theoretical results on graph Laplacian eigenvectors^36^. Consistent with this prediction, increasing the number of neurons further improved alignment (Supp. Fig. S5), although computational constraints limited exploration of this regime.

Altogether, these analyses suggest that strong local connectivity is essential to the emergence of geometric gradients, and that extending connections over larger spatial scales quickly degrades the alignment between dynamics and geometry.

### Geometric gradients are robust to randomized long-range connections

Neverthless, brain networks are characterized by the presence of numerous long-range connections, which markedly enhance routing efficiency and support the integration of functionally diverse processes across distant brain regions^22^. Although the EDR rule technically permits long-range connections, they are very scarce unless the characteristic length 1*/γ* is very large. As a result, most purely geometric models of connectivity fail to reproduce this key aspect of real brain networks^37^. More broadly, deviations from idealized wiring rules are expected in biological systems due to developmental variability, noise, and adaptive plasticity, irrespective of connection length.

To jointly account for long-range connections and stochasticity, we performed random double-edge swaps to the two network models considered so far. This procedure progressively and randomly introduced long-range connections while preserving each node’s degree and therefore the overall network density. To ensure a monotonous increase in the average connection distance ⟨*d*_*ij*_⟩ between nodes as edges are swapped, we prevented swaps that would produce edges shorter than the initial average connection length. Similar to the effect of gradually increasing 1*/γ*, increasing the fraction of swapped edges *ρ*_swaps_ led to a reduction in the average eigenmode-gradient correlation |*r*| for EDR networks initially parameterized at the optimal 1*/γ* = 0.0364 (Fig. 2**e**). Interestingly, we observed a slower decay of eigenmode-gradient correlations, with gradients remaining correlated to eigenmodes even with 10-20% of randomized edges. This effect was even more pronounced in networks initially wired using the connectivity radius rule at *h* = 0.12. For this case, eigenmode-gradient correlations remained high even after approximately half of the edges had been randomized (Fig. 2**f**). Thus, geometric gradients are tolerant to perturbations in connectivity.

To directly compare the effects of edge swapping with parameter manipulations in the EDR and radius models, we plotted the average eigenmode-gradient correlation |*r*| as a function of the mean connection length under all four connectivity rules (Fig. 2**g**). At comparable connection lengths, networks obtained by edge swapping from initially local architectures retained markedly higher eigenmode–gradient correlations than networks obtained by increasing either 1*/γ* or *h*. This contrast was particularly pronounced in the radius model: increasing *h* led to an abrupt breakdown of alignment with geometry, whereas introducing long-range connections via edge swaps caused a comparatively smaller degradation. The EDR rule underwent a similar degradation, although less pronounced since increasing 1*/γ* preserved non-negligible correlations over a broader range than for the hard-threshold radius model. Hence, the main limiting factor for the emergence of geometric gradients appears to be the extent of local connections and their intensity, since networks combining both local connectivity and sparse long-range connections align considerably better with geometry than those with a larger connectivity kernel.

Altogether, these results demonstrate that, while a backbone of local connectivity is necessary for the emergence of geometric gradients, the eigenmode-gradient correspondence is robust to the presence of randomized long range connections. As the average spatial extent of connections increases, however, this correspondence ultimately breaks down, regardless of the specific network topology. Importantly, these findings suggest that real neuronal networks, despite their substantial heterogeneity and abundance of long-range connections, may still exhibit geometric gradients, provided that the majority of connections remain spatially localized.

### Small wavelength eigenmodes progressively disappear as the connectivity radius increases

In the previous analyses, we quantified the alignment between function and geometry by averaging across multiple mode pairs, thereby obscuring potential mode-specific effects along the eigenspectrum. We therefore next examined correlations between individual eigenmodes and their corresponding functional gradients across different network parameterizations.

By visual inspection, progressively increasing the characteristic length 1*/γ* in EDR networks selectively eliminated correlations with small-wavelength modes (Fig. 3**a**). In contrast, the first long-wavelength modes were resilient to this change. At sufficiently large characteristic lengths, only the first geometric gradient persisted (Fig. 3**a**, right). To better visualize the effect of the connectivity kernel on individual eigenmode-gradient correlations, we plotted the diagonals of the mode correlation matrices at different exponential decay strengths (Fig. 3**b**). Strikingly, we observed abrupt cutoff points along rows (roughly indicated by arrows in Fig. 2**b**), suggesting that each geometric gradient exists only under a critical value of 1*/γ*. To detect these putative cutoff points, we identified the value of 1*/γ* corresponding to the maximal negative slope of |*r*_*i*_|(1*/γ*) for each of the first 50 mode pairs (Methods). We then compared these values to the characteristic spatial wavelengths *λ*_*i*_ of the corresponding eigenmodes, estimated from their spatial variograms (Supp. Fig. S6, Methods). We observed a linear relationship between wavelengths and the 1*/γ* cutoff points (Fig. 3**c**, *R*^2^ = 0.858), indicating that the position of each cutoff point is wavelength-dependent. Thus, the characteristic length of connections sets an abrupt spatial resolution limit beyond which geometric gradients with small wavelengths cannot be observed. We replicated this analysis using the connectivity radius model, observing even sharper cutoff points when increasing the radius *h* (Fig. 3**d-e**), whose positions scaled linearly with eigenmode wavelength (Fig. 3**f**, *R*^2^ = 0.952). This phenomenon can be interpreted analogously to the Nyquist limit in signal processing, whereby high-frequency components become unrecoverable when the sampling rate is insufficient.

**Figure 3.**
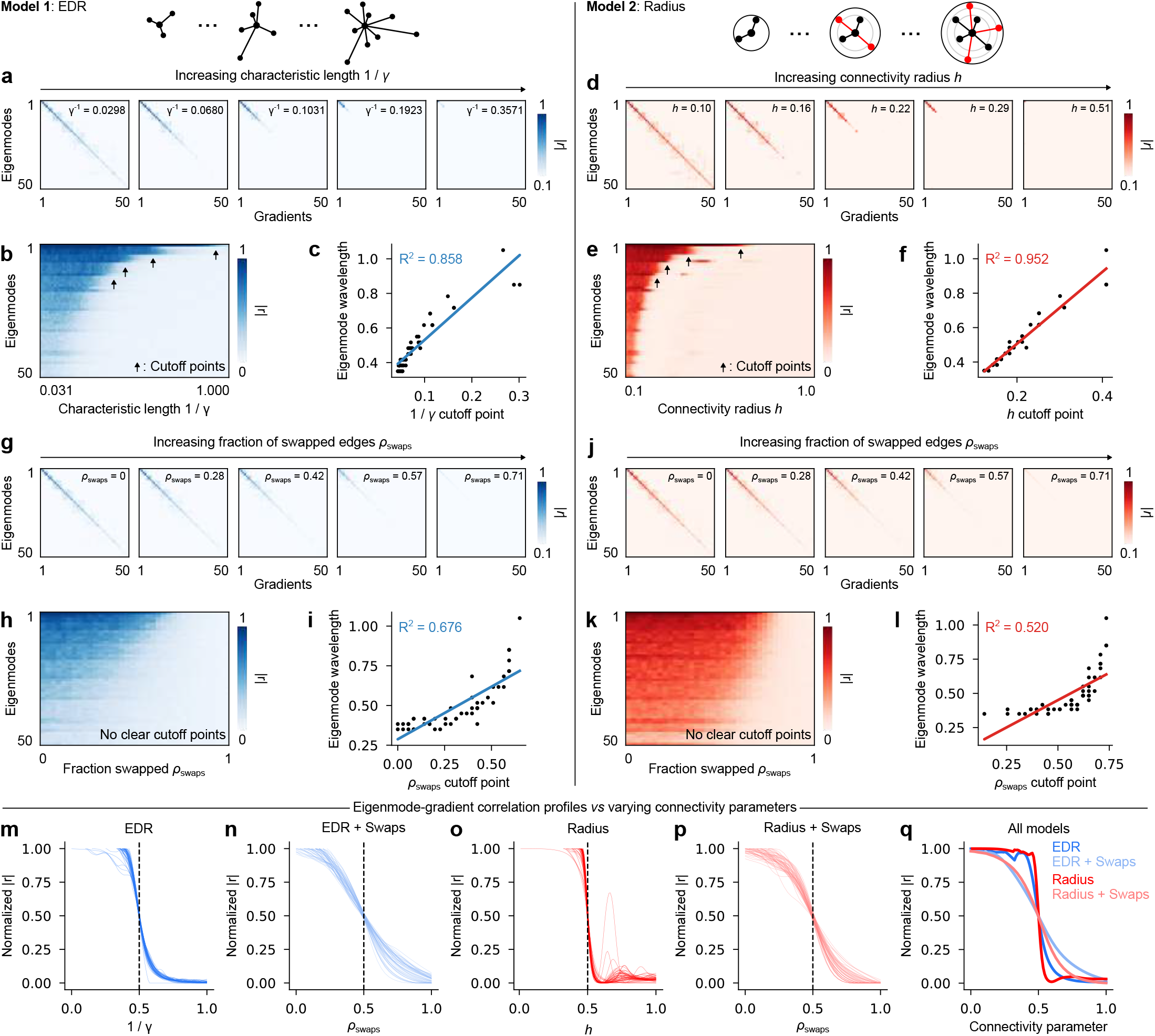
Degradation of small-wavelength modes with increasing long-range connections. (**a**) Five example eigenmode-gradient correlation matrices for increasing characteristic lengths 1*/γ*; each matrix is averaged over 10 different coordinate ensembles within the ellipsoid (*N* = 2500 neurons, 100 averaged simulations per coordinate set). (**b**) Eigenmode-gradient correlation matrix diagonals (columns) for different 1*/γ* values; arrows denote locations where eigenmode-gradient correlations abruptly drop (putative cutoff points). (**c**) Linear relationship between estimated eigenmode wavelengths and their 1*/γ* cutoff points. (**d, e, f**) Similar plots as **a, b, c**, but for the connectivity radius model. (**g, h, i**) Similar plots as **a, b, c**, but for the EDR model with edge swaps. (**j, k, l**) Similar plots as **a, b, c**, but for the connectivity radius with edge swaps. (**m**) Normalized and horizontally aligned row profiles (single eigenmode correlations) from **b.** (**n**) Normalized and horizontally aligned row profiles from **e**. (**o**) Normalized and horizontally aligned row profiles from **h**. (**p**) Normalized and horizontally aligned row profiles from **k**. (**q**) Average curves from the four previous panels.

We next conducted a similar analysis following double edge swaps to introduce long-range connections. Starting from locally connected EDR networks (1*/γ* = 0.0364), increasing the fraction of swapped edges *ρ*_swaps_ led to a more uniform degradation of eigenmode–gradient correlations, with the diagonal structure of the correlation matrix remaining clearly visible even when 20% of edges were swapped (Fig. 3**g**). Plotting the matrix diagonals as a function of *ρ*_swaps_ revealed no clear cutoff points beyond which individual gradients abruptly disappeared (Fig. 3**h**). Instead, correlations for all eigenmode–gradient pairs decayed smoothly with increasing edge swaps. Consequently, no strong linear relationship was observed between maximal slope points and eigenmode wavelength (Fig. 3**f**, *R*^2^ = 0.676). Similar results were obtained when edge swaps were applied to networks generated under the connectivity radius rule (Fig. 3 **j-l**). These results suggest that edge swapping leads to a gradual distortion of all geometric gradients at once, rather than the abrupt disappearance of individual eigenmodes. To further compare all four connectivity models, we normalized and horizontally aligned the individual eigenmode-gradient correlation curves (Fig. 3**m-q**), revealing sharp correlation drops when varying *γ* or *h* (Fig. 3**m**,**o**), and much smoother sigmoidal correlation drops following edge swaps (Fig. 3**n**,**p**). Note that for the radius model, weak residual correlations were observed for a small number of modes beyond their cutoff points (visible in Fig. 3**e** or Fig. 3**o**), however, these were substantially reduced relative to baseline correlations and likely reflect spurious matches arising from the mode-matching procedure.

Altogether, these analyses show that the minimal wavelength of geometric gradients scales with the characteristic length of connections. As connections are allowed to extend further in space, geometric gradients with shorter wavelengths first disappear, while long-wavelength gradients remain largely preserved. This result implies that in real neuronal networks, where connections extend over non-negligible distances, there should be an observable cutoff point beyond which eigenmodes and gradients no longer correspond.

### Spatial filtering artificially induces geometric gradients

Our numerical experiments demonstrate that the correspondence between eigenmodes and gradients is driven by short-range connectivity. An important implication is that any processing step that selectively enhances local correlations may artificially strengthen this relationship. Spatial filtering, ubiquitous in MRI preprocessing pipelines, blends signals from neighboring voxels, thereby enhancing local correlations. Recent work has shown that spatial filtering can spuriously induce smooth functional gradients in fMRI data, which may be misinterpreted as genuine connectopies^29^. This bias becomes particularly pronounced at finer spatial scales, such as when estimating gradients within individual brain regions.

To evaluate the influence of spatial filtering on geometric gradients, we substituted the simulated activity of neurons within the ellipsoid with temporally uncorrelated Gaussian noise (Fig. 4**a**). We spatially filtered the noisy signals using a Gaussian kernel (Fig. 4**b**), combining uncorrelated signals from nearby neurons according to a single parameter *σ* dictating the width of the gaussian kernel (Methods). By design, spatial filtering with a small kernel size of *σ* = 0.05 introduced high artificial correlations between nearby neurons (Fig. 4**c**). We computed functional gradients from both unfiltered and filtered correlation matrices, revealing strikingly smooth gradients in the filtered case (Fig. 4**d**). These filtering-induced gradients were strongly correlated with the geometric eigenmodes of the ellipsoid (average |*r*| = 0.793 across 20 modes; |*r*| = 0.604 across 50 modes; Fig. 4**e**). Thus, geometric gradients can be artificially induced by spatial filtering, even in the complete absence of correlation between the raw signals.

**Figure 4.**
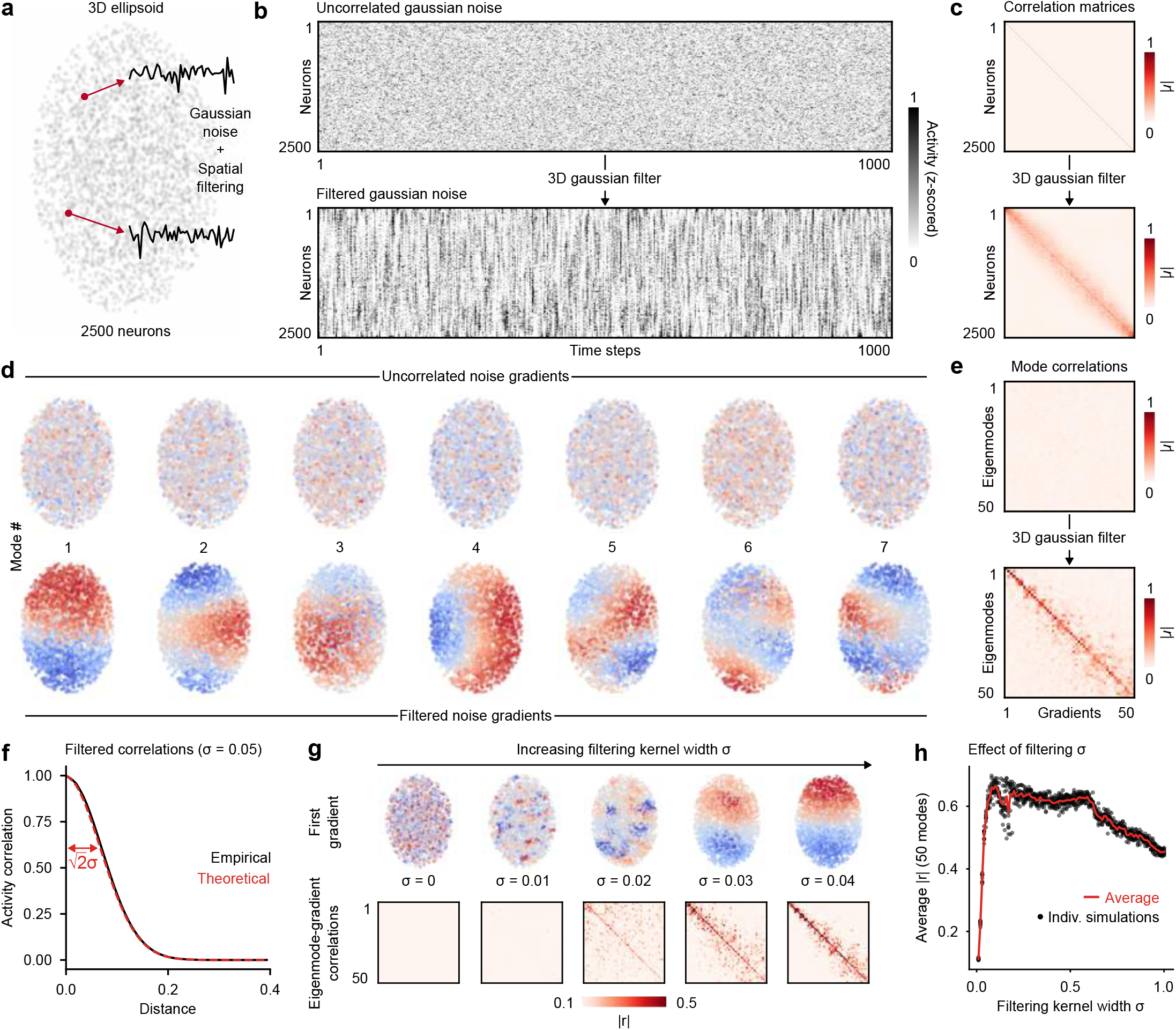
Spatial filtering of gaussian noise yields geometric gradients. (**a**) Schematization of the “simulations” for the noise filtering analysis; random gaussian noise signals are assigned 3D coordinates within an ellipsoid, then spatially filtered. (**b**) Top, matrix of random gaussian signals; bottom, same matrix after spatial filtering with *σ* = 0.05; neurons are ordered along the major ellipsoid axis. (**c**) Absolute correlations between noisy signals, before (top) and after (bottom) spatial filtering. (**d**) First 6 functional gradients before (top) and after (bottom) spatial filtering. (**e**) Eigenmode-gradient correlations before (top) and after (bottom) spatial filtering. (**f**) Distributions for theoretical (dashed red line) and empirical (black line) correlations between spatially filtered signals based on the distance between 3000 neurons in the ellipsoid geometry. (**g**) First gradient (top) and eigenmode-gradient correlation matrices (bottom) for increasing filtering kernel sizes *σ*. (**h**) Average eigenmode-gradient correlation |*r* | (over the first 50 mode pairs) for varying kernel sizes *σ*; black dots correspond to individual simulations (10 simulations per *σ* value, *N* = 1500 neurons), while the red curve is the average. Notice a notch at roughly *σ* = 0.2, which is likely an artifact of numerical discretization.

We next investigated the sensitivity of this effect to the width of the filtering kernel *σ*. First, we derived an analytical approximation for the filtering-induced correlations 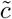 between two uncorrelated time series, encapsulated in the simple relationship 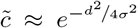 where *d* is the distance between the nodes (see Supplementary Mathematical Note). This relationship is exact in the limit of an infinite and homogeneous system. Crucially, the width of the filtering-induced correlation Gaussian is 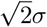, larger than the initial filtering kernel width. We validated this relationship numerically for small *σ* = 0.05 relative to the full size of the system to limit the influence of boundaries (Fig. 4**f**). This result suggests that spatial filtering induces correlations that deceptively extend beyond the spatial footprint of the kernel.

To further explore the implications on functional gradients, we evaluated the effect of *σ* on the average eigenmode-gradient correlation, observing the rapid emergence of smooth geometric gradients from very small filtering kernels (Fig. 4**g**, top). These smooth gradients were strongly correlated with geometric eigenmodes across all 50 mode pairs considered (Fig. 4**g**, bottom). In Fig. 4**h**, we plot the average eigenmode-gradient correlation | *r* | for *σ* ranging from 0 to 1, highlighting the very abrupt emergence of geometry with increasing *σ* from purely noisy, uncorrelated signals. Interestingly, eigenmode-gradient correlations remained high and decreased slightly for higher *σ* values that approached the characteristic length of the ellipsoid. This behavior contrasts with the effect of increasing 1*/γ* in the EDR model or increasing *h* in the radius model, as the eigenmode-gradient correspondence here persisted at large kernel sizes.

We also investigated the more realistic case of spatial filtering applied to a system that exhibits non-trivial correlations. In model 1, increasing the filtering kernel size morphed the local dynamics into broad activity patterns (Supp. Fig. S7**a**) and greatly enhanced local correlations between neurons (Supp. Fig. S7**b**). For sufficiently large filtering kernels (*σ* > 0.05), geometric gradients emerged, regardless of the network’s initial connectivity parameter (Supp. Fig. S7**c**). Interestingly, modest spatial filtering (*σ* < 0.05) in locally connected networks (1*/γ* ∼ 0.0364) decreased the eigenmode-gradient correlations (Supp. Fig. S7**c-e**), suggesting that filtering-induced correlations may, in some cases, interfere with raw correlations. Nevertheless, moving beyond this modest filtering regime eventually increased correlations above unfiltered levels.

These results underscore the high sensitivity of geometric gradients to spatial filtering, with artificial geometric features emerging from very small filtering kernels, consistent with previous reports on the confounding effects of spatial filtering in MRI studies^29^. Accordingly, spatial filtering may influence the interpretation of geometric features in imaging modalities that rely on such preprocessing steps, particularly in small brain structures.

### The larval zebrafish optic tectum exhibits geometric gradients

Our combined numerical analyses yield three testable hypotheses for experimental validation. First, homogeneous neuronal networks with predominantly local connectivity should give rise to geometric gradients. Second, neuronal arborizations that extend over non-negligible spatial scales should induce a detectable cutoff in the mapping between gradients and eigenmodes. Third, the characteristic wavelength of this cutoff point should quantitatively reflect the size of the network’s connectivity kernel.

To test these hypotheses, we used volumetric two-photon microscopy and calcium imaging to record spontaneous neuronal activity in the optic tectum of Tg(*elavl3*: H2B-GCaMP6s)^38^ zebrafish larvae at cellular resolution (Fig. 5**a**). The optic tectum is the main retinorecipient brain region of the zebrafish^39^ and exhibits spontaneous bursts of spatially compact neuronal assemblies^40;41^. This phenomenon is thought to arise from local recurrent excitatory connections among tectal interneurons^42^, a feature analogous to the spatially localized wiring implemented in our simulations.

**Figure 5.**
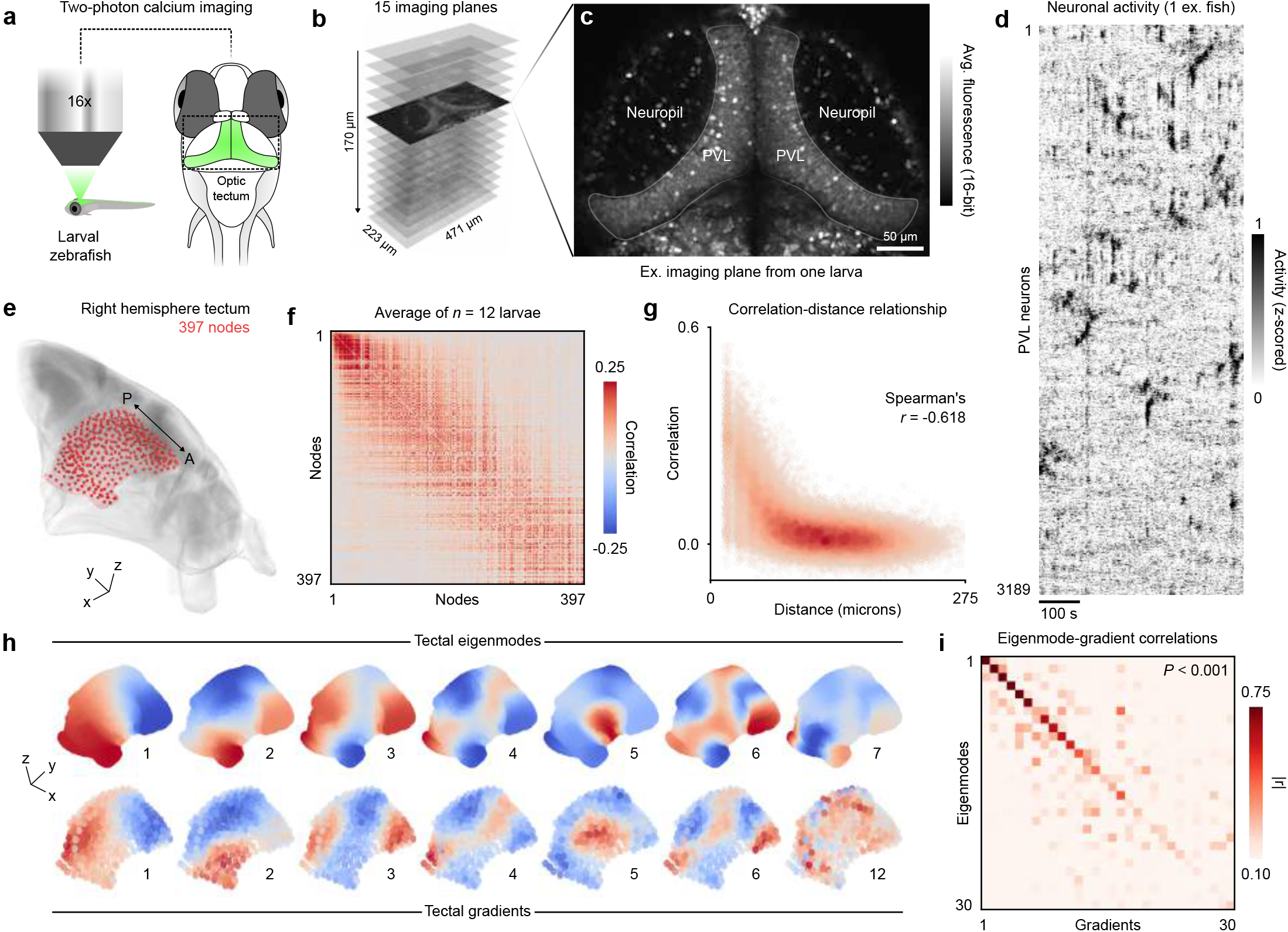
The larval zebrafish optic tectum exhibits geometric gradients. (**a**) Schematization of the microscopy experiment and field of view. (**b**) Dimensions of the multi-plane imaging volume. (**c**) One example time-averaged calcium imaging plane, highlighting the nuclear-localized calcium sensor, as well as the periventricular layer (PVL) and neuropil regions of the tectum. (**d**) Example time series of PVL neurons in the right hemisphere from one animal, sorted using the Rastermap^44^ algorithm to accentuate patterns. (**e**) 3D visualization and anatomical contextualization of the right tectum volume, with tectal nodes used to subdivide the region and allow inter-individual comparisons. (**f**) Pairwise correlations of node activity, averaged across 12 animals; the matrix is sorted along the antero-posterior brain axis. (**g**) Correlation-distance relationship obtained from the previous matrix, Spearman’s *r* = − 0.618. (**h**) Top, first 7 eigenmodes of the right tectum; bottom, corresponding functional gradients derived from calcium activity correlations; red and blue colors represent arbitrary positive and negative values, respectively. (**i**) Correlation matrix of tectal eigenmode-gradient pairs; the average correlation is globally significant when benchmarked against spatially shuffled gradient ensembles (*P* < 0.001, Methods).

We recorded 10 minutes of activity from thousands of neurons distributed throughout the entire tectal volume (15 imaging planes at ≈ 2 Hz, *n* = 12 larvae, Fig. 5**b**, Methods). We focused our analysis on cells within the periventricular layer (PVL neurons, Fig. 5**c**), a continuous and anatomically delineated structure. To avoid potential discontinuities across the midline, we restricted our analysis to the right hemisphere (2845 ± 578 neurons), although comparable results were obtained in the left hemisphere. Neuronal activity was characterized by low-amplitude calcium fluctuations punctuated by occasional large-amplitude transients involving tens to hundreds of neurons (Fig. 5**d**, Supp. Video 2). To average correlations across individuals, neuronal coordinates were registered to a common atlas space (mapZebrain atlas^43^, Methods). We then subdivided the right hemisphere tectum mask into 400 nodes of roughly equal size, to which we then assigned individual neurons (Fig. 5**e**, 397 nodes following exclusion criteria, Methods). We computed node time series by averaging single-neuron activity (7.16±4.35 neurons per node), then computed pairwise correlations between nodes (Fig. 5**f**). As previously reported^42^, we observed a strong distance-dependent decay in correlation strength within the region (Fig. 5**f**, Spearman *r* = −0.618), approaching baseline levels beyond approximately 100 *µ*m.

Next, we calculated tectal FC gradients from the group-averaged correlations and compared them to geometric eigenmodes derived from the 3D shape of the tectum (Fig. 5**h**, Methods). We observed positive correlations between eigenmodes and gradients across roughly the first 20 modes, beyond which the correspondence vanished (Fig. 5**i**). The statistical significance of this correspondence was assessed by comparing with null correlations obtained from spatially shuffled mode ensembles (*P* < 0.01, 1000 variogram-preserving permutation sets^32^, Methods). We ruled out spatial downsampling as a confounding factor, as eigenmode–gradient correlations were nearly identical across multiple coarse-graining levels (Supp. Fig. S9**a**). At the single-fish level, however, tectal correlations exhibited variability (Supp. Fig. S9**b**) and eigenmode-gradient correlations were weaker (Supp. Fig. S9**c**), likely reflecting limited statistical power due to the relatively short recording duration.

These results demonstrate that the intrinsic functional organization of the optic tectum is tightly coupled to its geometry, up to a cutoff point where the correspondence vanishes, thereby validating the first two hypotheses from our numerical simulations. Notably, these findings mirror observations made by Pang et al. in human subcortical structures^28^, extending them to a substantially smaller brain structure at cellular resolution in zebrafish larvae.

### The geometric cutoff point reflects tectal connectivity

The cutoff point observed in the eigenmode-gradient correspondence suggests that the connectivity kernel within the optic tectum is non-negligible relative to the size of the region. To test whether this cutoff point reflects the region’s underlying connectivity, we used a morphological dataset from the mapZebrain atlas^43^ to analyze the arborescence of 525 independently reconstructed tectal neurons (Fig. 6**a**). Unlike the radially extended connections used in our simulations, tectal interneurons extend both dendritic and axonal processes into an adjacent neuropil region^45^ (Fig. 6**a**). While a subset of neurons projects to the contralateral tectum or downstream hindbrain motor targets^46^, many interneuron arbors are confined entirely within the ipsilateral tectal neuropil (Fig. 6**b**, 18 example morphologies, top view).

**Figure 6.**
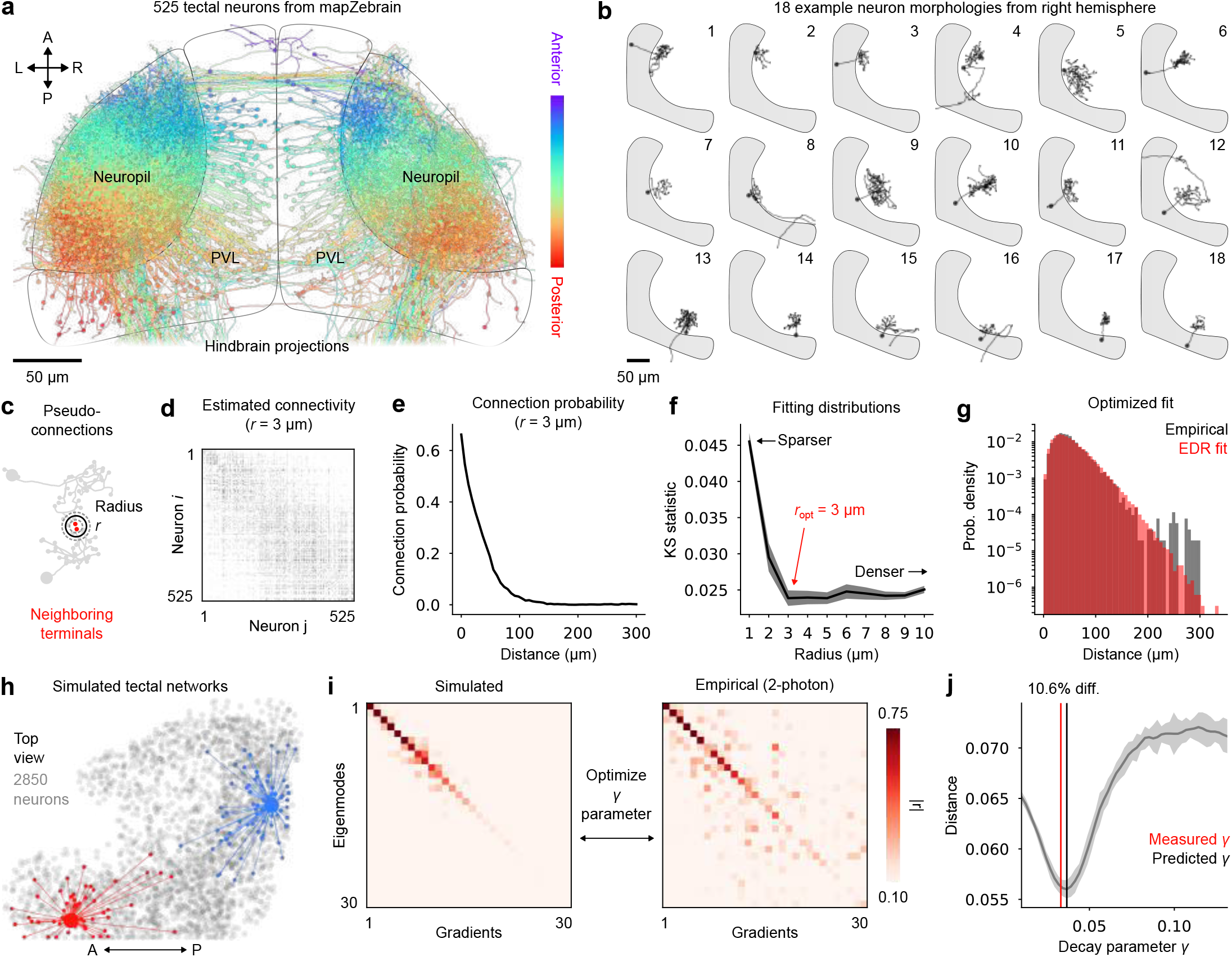
The geometric cutoff point reflects tectal connectivity. (**a**) Top view of 525 tectal neuron morphologies, colored by the antero-posterior location of the cell body, accentuating the topographic organization of neurites within the adjacent neuropil. (**b**) Example morphologies of 18 neurons, visualized from above. (**c**) Schematization of the connectivity estimation method; pseudo-connections are identified when neighboring terminals from different neurons are located within a certain radius *r* from one another. (**d**) Example binary connectivity matrix reconstructed using *r* = 3 *µ*m; neurons are sorted from anterior to posterior. (**e**) Connection probability as a function of distance, calculated from the previous matrix. (**f**) Kolmogorov-Smirnov statistic between empirical and EDR connectivity distributions (example in the next panel); an arrow indicates the optimal *r* value; the shaded region indicates standard deviation, 10 optimization runs per *r* value. (**g**) Empirical (gray) and EDR model (red) distributions of connection lengths at optimal *r* and *γ* values. (**h**) 2850 centroids in tectal geometry, visualized from above; example EDR connectivity profiles are highlighted in red and blue. (**i**) Left, average of 10 eigenmode-gradient correlation matrices obtained from simulations in tectal geometry using an optimized *γ* parameter (indicated in next panel); right, experimentally derived eigenmode-gradient correlation matrix (from Fig. 5**i**). (**j**) Euclidean distance between simulated and empirical eigenmode-gradient correlation matrices as a function of the *γ* parameter; the shaded region indicates standard deviation, 10 simulations per *γ* value; measured (red) and numerically predicted (black) *γ* values are indicated as vertical lines.

To estimate the region’s connectivity profile, we defined pseudo-connected neuron pairs as those whose neurite terminals lay within a distance threshold *r* of each other, drawing from the methodology established in a recent study^47^ (Fig. 6**c**). This simple method does not capture true synaptic connectivity, but rather uses arbor geometry to assess plausible synaptic contacts, favoring neurite endpoints where synapses are more likely^47^. Using this criterion, we reconstructed connectivity matrices across a range of distance thresholds *r* (Fig. 6**d**). Within a reasonable range of *r* values, the resulting pseudo-connectivity exhibited an exponentially decaying connection probability with distance, consistent with EDR connectivity (Fig. 6**e**, example shown for *r* = 3 *µ*m).

Because the *r* parameter dictates the density of the reconstructed network, we next sought an optimal threshold by comparing reconstructed connectivity to idealized EDR networks. Using soma coordinates, we generated EDR graphs across candidate *r* values and fit the empirical distribution of connection lengths by optimizing the decay parameter *γ* via a multi-step procedure (Methods). The Kolmogorov–Smirnov (KS) distance between empirical and model distributions was minimized at *r* = 3 *µ*m, for which an EDR model with *γ*_opt_ = 0.0331 (characteristic length 1*/γ*_opt_ = 45.5 *µ*m) closely matched the data (KS distance *D* = 0.022, Fig. 6**f–g**). To estimate uncertainty on *γ*, we simulated registration errors by perturbing soma and neurite coordinates by approximately 4 microns in random directions, reflecting typical error values in morphological atlases^43;48^ (Methods). Repeating the optimization process on 50 perturbed connectivity matrices yielded *γ*_opt_ = 0.0330 ± 0.0004 (mean ± standard deviation), suggesting this parameter is robust to small perturbations.

Next, we asked whether the previous eigenmode-gradient correlations in Fig. 5**i** could be reproduced by simulated dynamics on EDR networks, and whether the optimal *γ* would reflect the empirically estimated value. We first embedded *N* = 2850 neurons within a single tectal hemisphere to match the scale of our functional recordings, then simulated gradients resulting from EDR connectivity (Fig. 6**h**) while varying the *γ* parameter, as in the ellipsoid simulations. To match the cutoff point observed in the data, we minimized the Euclidean distance between simulated and empirical eigenmode-gradient correlation matrices (Fig. 6**i**). This distance exhibited a clear local minimum near *γ* = 0.0367, corresponding to a 10.6% deviation from the empirically measured *γ* = 0.0331 (Fig. 6**j**). Thus, the cutoff observed in calcium imaging data is accurately reproduced when the simulated connectivity has a characteristic length that mirrors morphological data. Although noise in the calcium imaging data could in principle influence the observed cutoff point by degrading FC gradients, we found the estimated cutoff wavelength to be robust across a wide range of simulated noise levels, up to three times the data standard deviation (Supp. Fig. S10).

Altogether, these results provide converging evidence from functional and structural imaging datasets in the zebrafish to support our third hypothesis: the geometric cutoff point in the eigenmode-gradient correspondence reflects a brain structure’s intrinsic spatial scale of connectivity.

### Whole-brain functional connectivity gradients do not reflect geometry

Our analyses thus far, together with prior neuroimaging results^28^, have focused on geometrically well-defined and relatively homogeneous systems such as brain regions or nuclei. However, brains are composed of multiple interconnected systems which, together, give rise to highly complex global dynamics^49^. Given the heterogeneous connectivity and prevalence of long-range projections, one should expect a weaker correspondence between eigenmodes and gradients at brain-wide scale.

To examine this question, we used our previous whole-brain two-photon calcium imaging dataset in zebrafish larvae^6^ to compute whole-brain FC gradients (*n* = 22 larvae, Methods). We first mapped individual neurons into 987 uniformly sized and non-overlapping nodes per brain hemisphere, then averaged single-neuron calcium fluorescence within each node before computing pairwise correlations. Correlations were then averaged across both hemispheres and animals (Fig. 7**b**, Methods). In contrast to the optic tectum, brain-wide functional connectivity exhibited only a weak decay with Euclidean distance (Spearman’s *r* = −0.298, Fig. 7**c**), indicating the presence of widespread long-range correlations.

**Figure 7.**
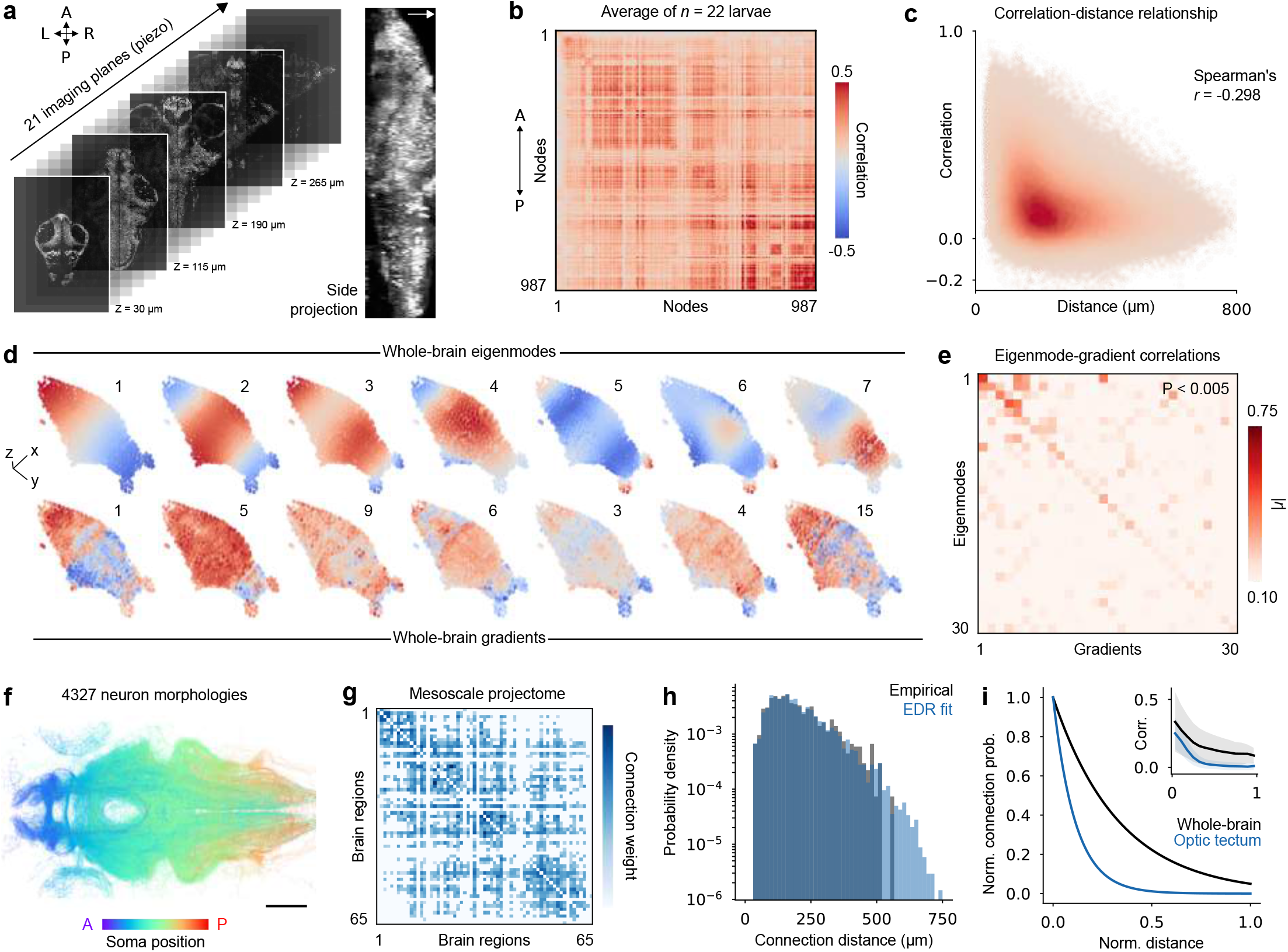
Whole-brain functional connectivity gradients do not reflect geometric eigenmodes. (**a**) Whole-brain imaging volume from one example animal; figure adapted from a previous study^6^. (**b**) Group-averaged whole-brain functional connectivity (*n* = 22 larvae, 5-7 days post-fertilization, 10 minutes of spontaneous activity); neurons are mapped in 987 nodes per hemisphere, and correlations are averaged across both hemispheres. (**c**) Correlation-distance relationship of whole-brain FC (Spearman’s *r* = − 0.298). (**d**) Top, whole-brain geometric eigenmodes; bottom, whole-brain functional connectivity gradients; both mode ensembles are projected onto 987 nodes per brain hemisphere and colors are scaled arbitrarily. (**e**) Whole-brain eigenmode-gradient correlation matrix; the correspondence is significant at the group level, that is, when eigenmode-gradient correlations are averaged over the matrix diagonal (*P* < 0.005, 1000 SA-preserving spatial permutations). (**f**) Neurite tracings from 4327 neurons, visualized from above; neurites are colored by the antero-posterior location of the cell body to accentuate certain projection patterns; scale bar, 100 *µ*m. (**g**) Mesoscopic connectivity matrix; connection weights are scaled arbitrarily. (**h**) Empirical (gray) and EDR model (blue) distributions of connection lengths at an optimal *γ* value. (**i**) Exponential connectivity profiles measured in the optic tectum (blue) and at whole-brain scale (black); spatial distances are normalized by the characteristic length of each system, thus a horizontal axis value of 1 corresponds to 270 microns or 750 microns in the tectum and the whole brain, respectively; inset, tectal and brain-wide correlation-distance profiles for direct comparison with structural exponential kernels; curves are average values while shaded regions represent standard deviation in each distance bin. Norm., normalized; prob., probability.

Next, we defined a single volumetric domain by segmenting the external brain boundary, treating the brain as a homogeneous structure without internal anatomical subdivisions (Methods). We computed single-hemisphere eigenmodes and compared them with corresponding functional gradients (Fig. 7**d**). We observed small, but nonetheless significant correlations between gradients and eigenmodes (Fig. 7**e**, *P* < 0.005, 1000 variogram-preserving spatial permutations, Methods). In particular, the main anteroposterior connectivity gradient correlated positively with the first eigenmode, but subsequent gradients were poorly matched (Fig. 7**e**). Thus, at the brain-wide scale, functional gradients are only weakly constrained by geometry.

A parsimonious explanation for this decoupling is that whole-brain connectivity violates the assumptions of local connectivity, due to the prevalence of long-range anatomical projections^27^. To test this hypothesis, we returned to the previous morphological dataset, which at the whole-brain level comprises 4327 neuron morphologies (Fig. 7**f**). Using these morphologies together with volumetric masks of 65 anatomical regions, we constructed a weighted mesoscopic projectome that captures inter-regional projection strengths (Fig. 7**g**, Methods). We next used the coordinates of brain region centroids to sample EDR graphs, then optimized the *γ* parameter as previously to match the empirical distribution of projection lengths. Brain-wide connection lengths were well captured by an exponential rule up to approximately 500 *µ*m, beyond which projections were largely absent (KS distance of 0.0207, Fig. 7**h**). The inferred characteristic length (1*/γ*_opt_ ≈ 250 *µ*m) corresponds to a broad connectivity kernel relative to brain size, with connection probability falling to 10% only at 575 *µ*m. To directly compare spatial scales, we normalized the tectal and brain-wide exponential rules by each system’s characteristic length (∼ 750 *µ*m for the whole brain; ∼ 270 *µ*m for the tectum; Fig. 7**i**). This comparison reveals a substantially narrower connectivity kernel within the tectum, consistent with its stronger geometry–function coupling. Rescaling the correlation profiles from Fig. S7**g** and Fig. 6**c** reveals a similar pattern at the functional level, with brain-wide correlations exhibiting considerably higher variability and remaining non-negligible over longer distances (Fig. 6*i*, inset plot). This further suggests that the EDR kernel size dictates the spatial spread of correlations.

Altogether, these results indicate that brain-wide networks are characterized by long-range pathways that support functional interactions between distant neural populations, thereby weakening the alignment between the brain’s global dynamics and its geometry.

### Spatial filtering in calcium imaging data

Lastly, we assessed how spatial filtering influences connectivity gradients derived from imaging data. We progressively applied spatial filtering to both tectal and brain-wide data, which drove high local correlations and resulted in a marked increase of the average eigenmode-gradient correlations (Supp. Fig. S11**a-b**). Peak correlations between eigenmodes and gradients were obtained at kernel widths of 50 and 18 microns for the whole-brain and the optic tectum respectively (Supp. Fig. S11**b**), representing around 7% of each system’s characteristic length (∼ 750 microns for the whole brain, and ∼ 270 microns for a single tectal hemisphere). Eigenmode-gradient correlations, averaged across the first 30 mode pairs, peaked at a lower value for whole-brain data (|*r*| = 0.436 versus |*r*| = 0.555 for the optic tectum). This behavior contrasts with our simulations, in which sufficiently strong spatial filtering ultimately drove strong eigenmode-gradient correlations, irrespective of the initial wiring topology (Supp. Fig. S7**c**,**d**). Moreover, the peak correlations in real data were observed at unreasonably large filtering kernel sizes (roughly 4 to 10 times the size of individual nuclei), suggesting geometric gradients are unlikely to emerge under conservative filtering regimes. The eigenmode-gradient cutoff wavelength observed in the optic tectum also remained mostly unchanged across the entire range of filtering strengths considered (Supp. Fig. S11**c**). These results indicate that, while spatial filtering could artificially drive geometric gradients in functional imaging data, more conservative filtering does not considerably increase eigenmode-gradient correlations, nor does it seem to bias other derived measures such as the geometric cutoff point.

## Discussion

The influence of geometry on nervous system organization is well established, with brain functions systematically arranged along the folded surface of the neocortex^7^. The prevalence of short-range connections in both small^43^ and large nervous systems^27^ supports spatially localized dynamics like bursting neuronal ensembles and traveling waves. Under certain modeling approximations, long-range connections may exert only a secondary influence to short-range interactions^24^. Such approximations imply that neural dynamics are fundamentally constrained by geometry in ways not fully captured by classical connectivity measures^25;28^. Whether connectivity or geometry provide greater explanatory power in shaping the brain’s macroscale dynamics has recently been debated^23;28;50;51^.

Here, we adopted a generative modeling approach to examine how geometry shapes emergent functional organization by simulating network dynamics across diverse spatial embeddings. Specifically, we investigated recent findings from Pang and colleagues who reported striking similarities between FC gradients and geometric eigenmodes in human subcortical structures^28^. To elucidate this alignment in different geometries, we imposed a singular constraint on simulations: that neurons are predominantly connected to other neurons that are nearby in space. Regardless of the precise wiring rule, dynamical model, or spatial embedding, this locality constraint consistently produced spatially structured correlations whose resulting connectivity gradients closely matched the geometric eigenmodes of the structure. This observation is consistent with theoretical results showing that, as network connectivity becomes increasingly local, geometry is imprinted in the graph itself^36;52^. In this regime, eigenvectors of the graph Laplacian, similar to diffusion-based FC gradients, converge toward those of the LBO defined on the graph’s spatial embedding^36^. We observed analogous spectral convergence across multiple gradient construction methods and, while the precise formal relationship between these operators remains unresolved, our results suggest that geometric gradients emerge naturally when local connectivity and correlations dominate. Importantly, by explicitly constraining structural connectivity through geometry, our modeling supports a view in which geometry and connectivity act jointly, rather than independently, to shape brain function.

While previous work has emphasized the role of wave-like dynamics in the alignment between geometry and function, our results show that geometric gradients arise from various network dynamics. Nevertheless, homogeneous wave-like dynamics are expected to yield geometric gradients due to the local correlations they induce^53^. The emergence of geometric gradients from various generative mechanisms and dynamical systems, however, highlights an important limitation of gradients derived from simple pairwise statistics: diverse dynamical processes can collapse onto nearly identical spatial patterns. This degeneracy is compounded by the arbitrary choice of pairwise interaction metric in neuroimaging studies^54^. Diffusion maps applied to alternative pairwise interaction matrices may yield spatial patterns that diverge from geometry, particularly for measures that are less coupled to spatial distance^54^. Given the limitations of linear correlations, together with the restricted spatial resolution of MRI, it remains unclear whether the geometric gradients reported by Pang and colleagues^28^ in the hippocampus, striatum and thalamus reflect wave dynamics, highly localized population bursts, or filtering effects. Traveling waves have been observed in the hippocampus using electrophysiological approaches^55;56^, and cortical waves can be decoded from subcortical activity in striatum and thalamus using Neuropixels probes^16^. Therefore, all three regions could plausibly support topographically organized activity patterns that translate into geometric functional gradients in macroscopic observations. While spatial filtering can accentuate these patterns, we found its influence to be modest in calcium imaging data when conservative filtering kernel sizes were used, particularly when the system did not exhibit geometric gradients in the first place. More broadly, our study provides an example of how temporally static statistical patterns such as FC gradients are inherently agnostic to the underlying dynamical processes, and should therefore be interpreted in conjunction with complementary analyses to more reliably infer their dynamical origins.

Our simulations further reveal that geometric gradients are robust to heterogeneities and long-range connections. Even when 10–20% of network connections were randomized, the correspondence between functional gradients and geometric eigenmodes remained largely intact, suggesting that this phenomenon is robust to noisy biological conditions. Although theoretical results predict exact convergence only in the limit of infinitesimally local connectivity^36^, we find that deviations from this idealized regime retain substantial eigenmode–gradient correlations in the low-frequency eigenspectrum. As shown in our simulations, spatially extended connectivity causes the geometric mapping between eigenmodes and gradients to break beyond a specific eigenmode wavelength that quantitatively reflects the size of the connectivity kernel. We experimentally tested this phenomenon using two-photon imaging in the larval zebrafish optic tectum, a predominantly locally connected structure that exhibits localized bursting activity^40^. As expected from these properties, we observed a good correspondence between eigenmodes and gradients in this region, as well as a clear geometric cutoff that could not be attributed to noise. Rather, this cutoff quantitatively reflected the strength of the exponential decay governing connection probabilities. While the optic tectum is not entirely homogeneous—featuring local connectivity variations and a layered cell-type organization^46;57^—its circuitry remains dominated by short-range connections consistent with its visuotopic organization^58;59^. Because complete electron microscopy reconstructions of the tectum are not yet available^60^, we estimated its connectivity rule using independently traced neuron morphologies and simple proximity rules. Although this approach likely biases the precise estimate of the decay parameter *γ*, the close agreement between the numerically inferred and empirically measured parameters indicates that the effective spatial scale of connectivity within the tectum was most likely well captured by both analyses. Together, our experimental results in zebrafish extend the findings of Pang and colleagues^28^ to a small vertebrate brain structure at cellular resolution, where activity signals are spatially resolved, non-overlapping, and unfiltered. This adds to recent studies highlighting larval zebrafish as a strategic model for comparative systems neuroscience^6;61^. Despite vast differences in brain size and complexity across species, core principles of brain structure and function often generalize and can be experimentally dissected in small model organisms^62^. Translating insights from human neuroimaging studies into experimentally tractable animal models is an essential step toward the validation of observations or theories established at macroscopic scales.

Although we demonstrate that the statistical phenomenon observed in subcortical nuclei can be reproduced in the larval zebrafish optic tectum, an important question remains: in what brain structures should we expect geometry and function to align? Our brain-wide analysis indicates that large-scale neuronal systems deviate from simple geometric rules, reflecting the prevalence of long-range connectivity and spatially decoupled activity. Similarly, in the human neocortex, the dominant FC gradient is spatially fragmented and closely tracks the default mode network^10^ (DMN), rather than the primary cortical eigenmode, which captures a smooth anteroposterior organization^28^. This decoupling could arise from preferential connectivity among distant cortical regions, or from structured subcortical inputs that impose non-geometric functional organization^63;64^. In general, we expect that the alignment between geometry and function is inherently scale-dependent. Isolated brain structures or nuclei may exhibit relatively homogeneous connectivity and dynamics, favoring geometric organization, whereas large-scale systems are assembled from multiple interacting subsystems with disparate wiring rules, breaking geometric assumptions^49;65^. Moreover, because geometric networks emerge from simple constraints, this phenomenon may also depend on developmental stage. Following the early establishment of local circuitry, long-range axonal tracts initiated by pioneer axons are reinforced through axonal growth or myelination^66^. In parallel, connections that are initially local can effectively become long-range as the system expands^67^. Such growth mechanism could support a progressive decoupling of neural dynamics from geometry as brain circuits mature and specialize. Consistent with this view, brain-wide correlation-distance relationships become weaker across development in humans^68;69^. Hence, geometric gradients are expected to emerge only under specific spatial and temporal conditions in functional imaging data. Nevertheless, deviations from geometric predictions may be equally informative by providing insights on the additional constraints that shape brain organization across scales and development^70^.

As our study relies on simple biophysical models with linear distance constraints, further work is required to clarify how geometry constrains synaptic connectivity—via factors such as wiring length^20^ or surface^71^ minimization, as well as volumetric occlusion^72;73^—and how this constrained wiring, in turn, imprints geometric features onto neuronal dynamics. From this perspective, geometry exerts its influence on brain activity through the structure–function relationship of brain networks, which serves as a critical intermediary^2;74^. Beyond chemical synapses, other biophysical processes such as extracellular neurotransmitter diffusion^4^ and gap junction-mediated synchronization^75^ also contribute to localized network dynamics. Exploring these alternative signaling mechanisms represents an important future direction to understand how brain function is grounded in physical space. As brain morphology varies across species, development, and disease^76^, understanding the influence of geometry on neuronal dynamics^28^ and brain organization^77^ is of both fundamental and translational interest. Our findings reinforce the central role of short-range interactions—a hallmark of brain networks and many other physical systems—as important drivers of geometric functional features in three-dimensional nervous systems.

## Methods

### Geometric eigenmodes

Geometric eigenmodes are the solutions to the Laplace-Beltrami eigenvalue problem defined on a Riemannian manifold, a type of geometric space where distance and curvature are locally defined. Mathematically, they are functions *ϕ*_*k*_ on that manifold, indexed by *k* ∈ ℕ, that satisfy Δ*ϕ*_*k*_ = *λ*_*k*_*ϕ*_*k*_, where Δ is the Laplace-Beltrami operator, *λ*_*k*_ are the corresponding eigenvalues, with *λ*_*k*_ increasing with *k*. The set of all *ϕ*_*k*_ forms an orthonormal basis of square-integrable functions over the manifold, ordered by increasing spatial frequency, i.e., lower-indexed modes correspond to smoother, slowly varying patterns, while higher-indexed modes exhibit finer spatial structure and more rapid oscillations. These eigenmodes represent intrinsic spatial harmonics of the geometric space and are commonly used to model patterns of diffusion or vibration constrained by geometry.

To derive the geometric eigenmodes of each geometry—that is, each three-dimensional Riemannian manifold—, we first generated volumetric tetrahedral mesh representations using the pygalmesh Python package. This process yielded approximately 30,000–40,000 tetrahedron vertices uniformly distributed throughout each volume and bounded within a unit cube (*x* ∈ [0, 1]), such that a distance of 1 represents a characteristic length scale common to all geometries considered. We then computed the geometric eigenmodes using a finite elements method (FEM) implemented in the LaPy package, solving for the first 100 eigenfunctions of the Laplace-Beltrami operator for each mesh, excluding the first constant eigenmode. The resulting eigenmodes are spatial functions defined over the mesh vertices, ordered by their eigenvalues (spatial wavelengths), with the first eigenmode corresponding to the smoothest pattern (longest wavelength) and higher-order eigenmodes representing increasingly fine-grained spatial variations (smallest wavelength).

### Wiring rules & simulations of neuronal activity

We simulated various dynamical models of neurons (described in the following section) embedded within 3D geometries. Although each model involved different parameters, all simulations followed a similar standardized protocol. Neuron coordinates were first randomly sampled from the set of vertices generated during the tetrahedral mesh construction described earlier. A binary connectivity matrix *W* ^bin^ was then generated using either exponentially decaying probabilities (EDR, model 1) or a deterministic connectivity radius (model 2). For the EDR model, neurons were connected with a probablity 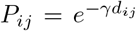 that decays with distance *d*_*ij*_ between neurons *i* and *j*. For the radius model, neurons were connected such that 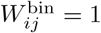 if *d*_*ij*_ < *h* and 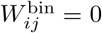 if *d*_*ij*_ ≥ *h*, where *h* is the connectivity radius parameters. Note that this binary network definition corresponds to a standard random geometric network model^78^. Next, *W* ^bin^ was transformed into a directed, weighted matrix *W* whose nonzero values were drawn from different distributions specific to each dynamical model, ensuring dynamics that are bounded, sustained, and exhibit rich, possibly chaotic behavior. Neuronal firing rates were then generated by numerically integrating the system over *T* = 2500 time steps using Euler’s method. To quantify emergent functional relationships between neurons, we computed an average correlation matrix *C* whose elements are the absolute pairwise Pearson correlation coefficients of neuronal firing rates averaged over 50 independent simulations. In each run, the connectivity weights and initial conditions were reinitialized, while the neuron position and the binary structure *W* ^bin^ remained fixed. This ensured a representative sampling of possible dynamics unfolding on the same geometric graph. We used absolute correlations to capture both large positive and negative relationships between neurons, though in practice, the largest-magnitude correlations were predominantly positive and similar results could be achieved without taking the absolute value, either by rescaling the matrix or clipping negative values. Finally, the resulting average correlation matrix *C* was used for subsequent gradient analysis. For some analyses, we repeated the entire simulation protocol 10 times per geometry and parameterization—each time resampling node coordinates, constructing a new connectivity graph, simulating dynamics, and averaging correlations—to reduce bias introduced by the randomized node coordinates and to suppress noise in the eigenmode-gradient correlations. All numerical simulations were implemented in PyTorch using CUDA, enabling efficient GPU parallelization and substantially accelerating the numerical integration of large dynamical systems.

### Dynamical models

#### Model #1: Chaotic firing rate network

For the first and main dynamical model used throughout most of the study, we simulated spontaneous firing rate dynamics—also known as the graded response model—from the following equations^79^:

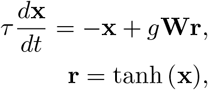

where **x**(*t*) ∈ ℝ^*N*^ describes an internal variable for each neuron at time *t*, analogous to a membrane potential, while **r**(*t*) is the vector of firing rates at time *t* obtained through a nonlinearity tanh(**x**), with tanh applied element-wise. The parameter *τ* dictates the time constant of neurons, and *g* is the network coupling strength, which brings the dynamics into a chaotic regime for *g* > 1 in the thermodynamic limit^31^. The connectivity matrix *W* ∈ ℝ^*N ×N*^ has entries drawn independently and identically from a normal distribution 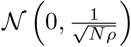, with positive and negative values interpreted as excitatory and inhibitory synaptic weights, respectively. The parameter *ρ* reflects the empirical average connection density, whose exact value fluctuates with node coordinates and connectivity kernel size (networks with small 1*/γ* and small *h* are sparser, and vice-versa). Crucially, the *ρ* normalization of connection weights ensures that the overall strength of inputs to each neuron remains constant as the connectivity radius *h* increases, preventing runaway activity in denser network configurations (Supp. Fig.S4). We used a stronger coupling of *g* = 3 to compensate for network sparsity and to ensure rich chaotic activity time series (Fig.1 **c**). We used a small time constant of *τ* = 3 to produce faster dynamics and ensure stabilized correlations over the full integration time. Initial conditions **x**(0) were sampled randomly and uniformly in the range [−1, 1] before numerically integrating the system.

#### Model #2: Chaotic firing rate network + Dale’s law

For the second dynamical model, we adapted the first dynamical model by changing the neuronal activation function and applying Dale’s law to the connectivity matrix. Instead of the standard **r** = tanh(**x**), which varies between 1 and 1, we used a strictly positive activation function defined in a previous study^33^,

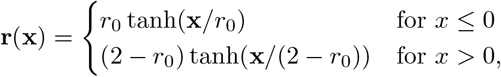

with firing rates ranging **r**(*t*) from 0 to 2. The *r*_0_ parameter represents a fixed background firing rate, which we set to *r*_0_ = 0.1. Note that for *r*_0_ = 1, this function reduces to the standard tanh function. In addition to this change in the neuron dynamics, we applied Dale’s law, that is, we imposed strictly positive or negative signs on the columns of *W*. This ensures that each neuron’s outputs are strictly excitatory or inbibitory, respectively. We used a 1: 1 ratio of excitatory and inhibitory neurons to maintain properly balanced oscillatory dynamics. These combined changes to the network architecture led to qualitatively different activity, with more spatially patterned population fluctuations that were not present in the more chaotic dynamics of model #1.

#### Model #3: Kuramoto-Sakaguchi oscillators

For the third dynamical model, we considered a network of *N* coupled Kuramoto-Sakaguchi oscillators, each interpreted as a neuron, such that the phase *θ*_*i*_ of neuron *i* evolves according to^34^

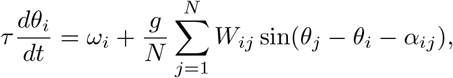

where *τ* is a global time constant, *ω*_*i*_ is the natural frequency of neuron *i, W* is the connectivity matrix, *g* is a global coupling strength parameter, and *α*_*ij*_ is a phase-lag parameter. For this model, the weights in *W* remained binary, with *W*_*ij*_ = 1 indicating the presence of a connection from neuron *j* to neuron *i*, and *W*_*ij*_ = 0 otherwise. The phase lag matrix *α* was obtained by multiplying the euclidean distance matrix by an arbitrary scaling parameter. This distance-dependent phase lag approximates propagation delays along structural connections of variable lengths^80^ and aims to maintain the network in a chaotic regime. We set *g* = 4000, *τ* = 20, and we randomly sampled natural frequencies according to *ω* ∼ 𝒩 (2, 0.1). Phases were initialized randomly and uniformly in the [0, 2*π*] range. Parameter values were selected by a linear sweep to maximize the alignment between geometric eigenmodes and FC gradients. Interestingly, chaotic dynamics emerged regardless of the phase lag scaling parameter in EDR networks. For this reason, we set *α* = 0, thus reducing to simple Kuramoto dynamics^81^. For radius-based networks, however, an optimal scaling parameter of 50 was required to yield chaotic dynamics, hence why we refer to the expanded KS model.

#### Model #4: Binary stochastic neurons

For the fourth dynamical model, we implemented a simple binary stochastic neuronal network where the *N* -dimensional network state vector **x** is binary, with ones representing active neurons and zeros representing inactive neurons. This model can be viewed as a Cowan-type Markov chain model of neurons^82;83^ with a clipped ReLU activation function. More explicitly, the state evolution is governed by two parameters: *P*_*a*_, the probability that a neuron becomes active per active presynaptic neuron, and *P*_*d*_, the probability that a neuron becomes inactive spontaneously at each integration step. For this model, connectivity matrix *W* remained binary, and Dale’s law was applied in a 1: 1 ratio to ensure balanced excitatory and inhibitory interactions. At each time step, inputs from active neurons were summed at each postsynaptic neuron, then the resulting integer number of inputs was multiplied by *P*_*a*_ to yield an activation probability, which we clipped in the range [0, 1]. Importantly, for a sufficient number of positive inputs, postsynaptic neurons are guaranteed to become active. We initialized the network with roughly 10% of active neurons, and used *P*_*a*_ = 0.18 and *P*_*d*_ = 0.4 to obtain sustained chaotic dynamics.

#### Model #5: Spiking neurons + calcium dynamics

For the fifth and final dynamical model, we implemented a discrete-time version of a conductance-based leaky integrate-and-fire (LIF) neuronal network^35^ with simplified calcium dynamics. The network consists of *N*_*E*_ excitatory neurons and *N*_*I*_ inhibitory neurons, whose membrane potentials and synaptic conductances evolve according to

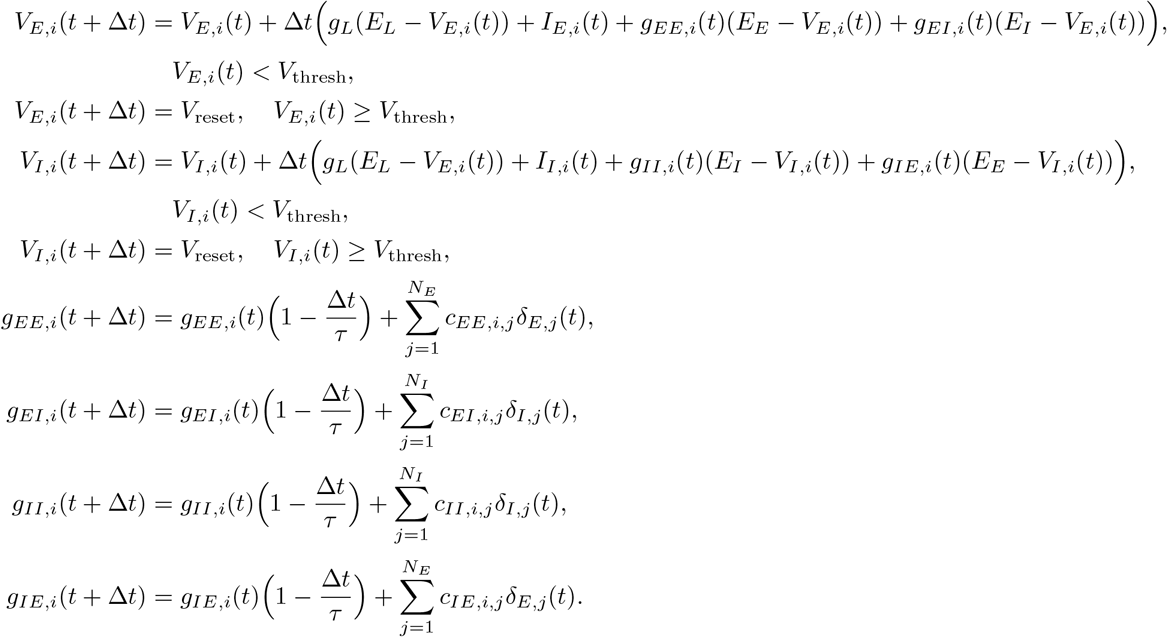

**V**_*E*_(*t*) and **V**_*I*_ (*t*) represent the membrane potentials of excitatory and inhibitory neurons, respectively, while **g**_*EE*_ (*t*), **g**_*EI*_ (*t*), **g**_*IE*_ (*t*) and **g**_*II*_ (*t*) represent the conductances from *E* → *E, I* → *E, E* → *I* and *I* → *I* populations, respectively. The terms *δ*_*E,i*_(*t*) and *δ*_*I,i*_(*t*) are equal to 1 if excitatory or inhibitory neuron *i* emit an action potential at time *t*, else they equal 0. Neurons spike once their membrane voltage exceeds a threshold *V*_thresh_, upon which the voltage is immediately reset to *V*_reset_ the following time step. *I*_*E*_ and *I*_*I*_ are constant inputs delivered to the excitatory and inhibitory populations. *E*_*L*_, *E*_*E*_ and *E*_*I*_ are the leak, excitatory, and inhibitory reverse potentials, and *g*_*L*_ is the leak conductance. Finally, *τ* is the synaptic time constant, and Δ*t* is the integrator time step. To simulate the model, we distributed excitatory and inhibitory neurons uniformly throughout the volume, connected them with binary synaptic weights, then used the following set of parameters to generate sustained chaotic dynamics: *g*_*L*_ = 0.15, *E*_*L*_ = −75, *E*_*E*_ = −40, *E*_*I*_ = −90, *V*_thresh_ = −55, *V*_reset_ = − 75, *τ* = 0.1, *I*_*E*_ = 5, *I*_*I*_ = 0, and Δ*t* = 0.025. We finally convolved spike trains with an exponential kernel (*τ* = 10) to emulate intracellular calcium dynamics and permit correlations between the sparse spiking vectors.

### Connectivity gradients and diffusion maps

Connectivity gradients are low-dimensional representations of the large-scale organization of a network, revealing principal axes of variation in connectivity. When applied to brain data, they capture how neural units are functionally or structurally organized along continuous topographies.^10^ Neural units with similar values along specific gradients tend to exhibit similar connectivity patterns, i.e., having several neighbors in common. Connectivity gradients were calculated using the diffusion maps algorithm^30^. This standard approach consists of calculating the eigendecomposition of a diffusion operator *P*_*α*_ depending on a parameter *α* ∈ [0, 1], and defined as

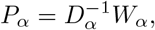

with *W*_*α*_ = *D*^−*α*^*CD*^−*α*^, *C* being the correlation matrix under study, *D* is a diagonal matrix with entries 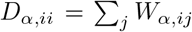 and *D*_*α*_ is a diagonal matrix with entries *D*_*α,ii*_ = ∑_*j*_ *W*_*α,ij*_. The diffusion operator *P*_*α*_ is row-stochastic and therefore has eigenvalues satisfying 1 = *λ*_0_ > *λ*_1_ ≥ *λ*_2_ ≥ · · ·. The connectivity gradients *G*_*i*_ correspond to the leading non-trivial eigenvectors *ψ*_*i*_ of *P*_*α*_, excluding the trivial eigenvector associated with *λ*_0_ = 1. In diffusion maps, these eigenvectors are rescaled by their eigenvalues according to

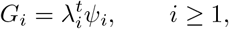

where *t* is the diffusion time. The value of *α* dictates how strongly the density of the sample points (or nodes if C is derived from a graph) on the underlying manifold is taken into account when computing the gradients. Importantly, for specific values of *α*, this family of operators reduces to known diffusion operators. For *α* = 1, *P*_*α*_ approximates the Laplace-Beltrami operator on the manifold, removing all influence of the density. For *α* =^1^*/*2, *P*_*α*_ approximates Fokker-Planck diffusion. Finally, for *α* = 0, *P*_*α*_ reduces to the transition matrix for the random walk on *C, D*^−1^*C*. For most of our analyses, we used *α* = 0.5, which is standard in the gradient literature. For numerical calculations, we used the BrainSpace implementation of diffusion maps^84^, which uses the following normalization of the eigenvalues, 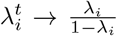 thus eliminating the additional choice of parameter *t*. Note that in this study, gradients are not used to embed nodes. Rather, we correlate them to eigenmodes individually, such that the diffusion time parameter is irrelevant. Moreover, the density of sample points is approximately uniform for both our numerical simulations and real parcellated data, hence it is expected that *α* would have almost no influence on the results.

In Supp. Fig.S3, we compared three gradient derivation methods. We first demonstrated that geometric eigenmodes can be obtained across the full range of the *α* parameter. Alternatively, we computed gradients from a slightly different matrix. Instead of using the raw correlation matrix *C*, we computed the pairwise correlations of connectivity profiles (rows of *C*), then proceeded with the diffusion maps on this resulting matrix. As a third method, we mimicked the connectopic mapping technique^85^ used in Pang et al^28^, where gradients within a brain region reflect similarities in connectivity profiles with other brain regions. To do this, we simulated two separate neuronal populations of size *N* and *M*, with the first one sending random inputs to the second one. Then, we generated an *N* × *M* correlation matrix of firing rates across the two systems, and computed a *N* × *N* matrix of connectivity profile similarity for nodes within the first system, relative to the second one. We finally used this similarity matrix to compute gradients using diffusion maps.

### Mapping eigenmodes and gradients

To compare eigenmodes and gradients, we computed their pairwise absolute correlations *R*_*kℓ*_ between the values of eigenmode *k* and those of gradient *ℓ*, then solved the following linear sum assignment problem max ∑*R*_*kℓ*_*X*_*kℓ*_, where *X* is a boolean matrix where *X*_*kℓ*_ = 1 if and only if eigenmode *k* is assigned to gradient *ℓ*. This procedure is analogous to reordering the columns of *R* to maximize the values on the diagonal. We used the scipy implementation^86^, which uses a modified Jonker-Volgenant algorithm^87^.

### Double edge swapping procedure

The goal of double edge swaps was to preserve the node degrees of geometric graphs while randomizing their connections and introducing long-range connectivity. To do so, we began by selecting two edges at random, say (*u*_1_, *u*_2_) and (*v*_1_, *v*_2_). We then swapped those edges, by replacing them with the new edges (*u*_1_, *v*_2_) and (*v*_1_, *u*_2_). The weights of the initial edges were assigned randomly to the new edges to preserve the total weights (and thus the average weighted degree, but not the individual weighted degrees) of the graph. To ensure that the edge swapping procedure monotonically increases the connectivity distance on average, we rejected swaps where any of the new edges would have a length less than the average connection length in the EDR model, or less than the connectivity radius *h* in the second model. We repeated this process by selecting two random edges of length smaller than *h* and swapped them until a certain fraction *ρ*_swaps_ of the network’s edges had been swapped.

### Identifying geometric eigenmode cutoff points

To locate geometric cutoff points, we identified the inflexion point in the sigmoidal profiles visible in Fig.3, corresponding to the point where a given eigenmode is abruptly disappearing from connectivity gradients. To do so, for each eigenmode, we computed the correlation |*r*| with its corresponding gradient for each characteristic length 1*/γ* or connectivity radius *h*, and determined the interval [*h*_0_, *h*_1_] for which *r* decreases the most, and for which the adjacent intervals also present a decrease in |*r*|. This last condition serves to guarantee that the cutoff point is in the proper region of the curve and to protect against noise effects. The cutoff value of 1*/γ* or *h* was then determined as the middle point of the maximally decreasing interval, 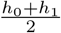. Overall, this method treats each eigenmode separately to identify a relationship between the size of the connectivity kernel and eigenmode wavelengths.

For experimental data, we used a continuous piecewise linear fit on the eigenmode-gradient correlation matrices to identify a “cutoff wavelength”, that is, an inflexion point where correlations reach baseline levels. This piecewise linear fit consists of a decreasing linear part, − *a*# + *b*_1_ (where # refers to eigenmode number), and then a constant (horizontal) part, *b*_2_. We found the set of paramaters *a, b*_1_, *b*_2_ and #_*cutoff*_ that maximizes the *R*^2^ of the regression, and identified the rounded optimal *λ*_*cutoff*_ as the wavelength of the eigenmode corresponding to the cutoff point.

### Estimating eigenmode wavelengths

To estimate the eigenmode wavelengths in arbitrarily shaped geometries, we computed the variogram of each eigenmode *ϕ*_*k*_. For each eigenmode, we randomly selected 2500 vertices and calculated the pairwise squared differences of their eigenmode values, 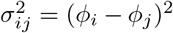. These distances were then averaged within 30 spatial distance bins ranging from 0 to This subsampling process was repeated 10 times per eigenmode to generate an average variogram. The half-wavelength of each eigenmode was estimated as the distance *d* at which the first peak of the variogram occurred, corresponding to the shortest average distance between regions of maximally opposing signs in the spatial function. We multiplied these distances by 2 to recover full approximate wavelengths.

### Spatial filtering

In both simulations and real data, spatial filtering was applied by combining node time series according to the following relationship:

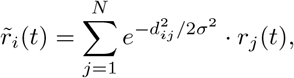

where *r*(*t*) are the raw noisy signals, *σ* is the Gaussian kernel width, and *d*_*ij*_ is the distance separating neurons *i* and *j*. This corresponds to a standard Gaussian filter applied to the continuous set of network coordinates.

### Animal model

All microscopy experiments were conducted *in vivo* on 5-7 dpf Tg(*elavl3*: H2B-GCaMP6s)^38^ zebrafish larvae. Larvae were raised in embryo medium (13.7mM NaCl, 0.54mM KCl, 1.0mM MgSO_4_, 1.3mM CaCl_2_, 0.025mM Na_2_HPO_4_, 0.044mM KH_2_PO_4_, 4.2mM NaHCO_3_, pH 7.2 with HCl 12N) at a density of 1 larva per mL in Petri dishes placed in an incubator at 28°C on a 14:10 hour day/night cycle. From 5 dpf onward, the medium was replaced daily and larvae were fed live *Tetrahymena thermophila* CU428.2 (Cornell University). *T. Thermophila* were grown in Petri dishes in sterile SPP medium (2% proteose peptone, 0.1% yeast extract, 0.2%glucose, 33 *µ*M FeCl_3_, 250 *µ*g/mL Penicillin/streptomycin, 0.25*µ*g/mL Amphotericin B) at room temperature and harvested in the stationary phase. They were then washed from their SPP medium using three centrifugation steps (2min at 0.8g each), followed by pellet resuspension in embryo medium. All protocols were approved by the animal care committee of Université Laval.

### Two-photon calcium imaging

Larvae were embedded in 2% low-melting point agarose (Invitrogen 16520100) in a 30 mm glass bottom petri dish (Mattek P35G-1.0-14-C) and positioned upright for brain imaging using a toothbrush bristle without direct contact with the animal. The preparation was submerged in embryo medium and placed under the microscope for imaging. All imaging experiments were conducted on a Scientifica SliceScope resonant two-photon microscope (SciScan LabView software) equipped with a Spectra Physics Insight X3 tunable laser. Fluorescence was collected using a piezo-driven 16x water-dipping objective (Nikon, 0.8 NA), reflected by a T585lpxr dichroic mirror, and bandpass filtered (505/119nm) before detection with a GaAsP detector. High-resolution anatomical stacks centered on the tectum were first collected across the full depth of the brain (1024 × 494 pixels, ≈ 200 planes, 2 microns in z-spacing, 24x frame averaging, zoom level 1.1) using an excitation wavelength of 860 nm, which is near the isosbestic point of the calcium indicator. Following the structural scan, calcium imaging of the tectum was performed at an excitation wavelength of 920 nm. We sampled neuronal activity from 16 imaging planes (1024 × 494 pixels, pixel size approximately 0.46 ×0.46 *µ*m, zoom level 1.25, 2 Hz volume rate) spaced approximately 12 microns apart, and we excluded the first plane due to piezo recoil artifacts induced by the sawtooth scanning pattern. We delimited the imaging volume by positioning the first plane at the top of the brain where labeled nuclei are first encountered, and then positioned the last plane slightly below the tectum. Power measurements after the objective were maintained at 20 mW, and functional imaging experiments lasted 10 minutes under spontaneous conditions with static red light projected on a screen under the larvae. For the whole-brain imaging experiments, we used similar parameters which are detailed in a previous publication^6^. We analyzed the spontaneous portion of these experiments which lasted roughly 10 minutes, again with static red light projected under the larvae.

### Data processing

Calcium imaging planes were corrected for nonrigid lateral motion using the NoRMCorre algorithm^88^, which is implemented in Python within the CaImAn suite^89^. We used a large patch size of 240 × 240 pixels with a 120 pixel overlap to prevent local registration artifacts. Then, we segmented fluorescent nuclei from the time-averaged calcium imaging frames of each plane using Cellpose 2.0^90^. To properly detect the densely labeled nuclei, we trained a custom model on 10 different patches containing roughly 100 neurons each using the expert-in-the-loop method. We then extracted fluorescence signals by averaging pixels within each region of interest (ROI), and detrended the signals using a custom method described in a previous study^6^. Briefly, we used a minimum filter to identify the time-varying baseline of each signal (temporal window of 60s), then normalized the signals by their baselines to obtain relative fluorescence measurements, Δ*F* (*t*) = (*F* (*t*) − *F*_0_(*t*))*/F*_0_(*t*), where *F* (*t*) is the raw activity trace and *F*_0_(*t*) is the inferred baseline. To map each neuron in a standardized anatomical coordinate system, we used ANTs registration^91^ to register the anatomical volume acquired at *λ* = 860 nm to an atlas template brain generated in the same transgenic line (mapZebrain^43^), using standard alignment parameters^92^. To ensure a proper alignment of the tectal volume within the larger brain volume, we zeroed the reference template values located outside our approximate imaging field of view. We repeated this alignment procedure between the functional and anatomical imaging volumes acquired at *λ* = 920 nm and *λ* = 860 nm respectively, then used the antsApplyTransformsToPoints function twice to transfer the 3D coordinates of neurons from functional to anatomical stacks, then from anatomical to atlas coordinates.

### Computing tectal gradients and eigenmodes

To compute functional connectivity gradients in the optic tectum, we first identified neurons whose transformed coordinates were located within the tectal periventricular layer using a brain region mask defined in the atlas. Then, we subdivided the right hemisphere mask into 400 evenly spaced subregions using k-means clustering, and then assigned individual neurons to their nearest neighboring node. Importantly, neurons were assigned to one node only, thus avoiding the blending of signals and artificial strengthening of correlations between adjacent nodes. We averaged neuronal activity within each node, then correlated these signals to yield anatomically comparable correlation matrices across animals, which we then averaged before computing gradients on the group-average matrix, which exhibited very few negative values. We used absolute correlations to compute gradients, yielding similar results than thresholded correlations clipped in the range [0, 1] to remove negative values. For gradients computed on individual FC matrices (Supp. Fig.S9 **b**), negative values were more prominent, and we rescaled matrices in the range [0, 1] before computing gradients. For eigenmode calculation, we used the right hemisphere of the 3D PVL binary mask from the brain atlas as a volume for mesh generation. We generated a tetrahedral mesh and solved the eigenmodes using the FEM from LaPy as described earlier.

### Estimating tectal connectivity

To estimate the exponential connectivity kernel within the optic tectum, we leveraged a dataset of 4327 independently reconstructed neurons from mapZebrain, compiled across different animals^43^. Of these neurons, 525 have their soma located within the periventricular layer (PVL), corresponding to where nuclei are located in our calcium imaging dataset. For each PVL neuron, we identified arbor terminals located within the adjacent tectal neuropil as putative synapses, and then identified pseudo-connected partners, that is, other neurons with terminals located within *r* microns of the neuron’s terminals. To focus on within-hemisphere connectivity, we only counted pseudo-connections between neurons whose somas are both located within the same hemisphere. While synapses may be observed all along neurites in zebrafish^60^, their precise locations are uncertain in light microscopy datasets due to lack of protruding spines. Focusing on neurite endpoints therefore aims to reduce false positives.

### EDR model fitting

To test whether empirical connectivity in the tectum or at the whole-brain level follows an EDR, we compared empirical distributions of connection lengths (Fig.6 **g**, Fig.7 **h**) to distributions of edge lengths generated by an exponential distance rule (EDR) model applied to neuron or brain-region coordinates. We optimized *γ* by minimizing the Kolmogorov–Smirnov (KS) distance between empirical and model distance distributions using an iterative, multi-stage search procedure. Starting from an initial estimate *γ*_0_ = 0.05, we evaluated KS distances over 25 linearly spaced *γ* values within a symmetric window of width 0.05 centered on *γ*_0_. For each candidate *γ*, model distance distributions were generated by repeating the EDR sampling procedure five times and pooling the resulting edge lengths. The KS distance profile across the parameter sweep was smoothed using a Gaussian filter (*σ* = 2), and the *γ* value corresponding to the local minimum was selected as the current estimate *γ*_opt_. The search window was then narrowed by a factor of 0.66 and recentered on *γ*_opt_, and the procedure was repeated. This iterative refinement was performed for 10 iterations, yielding a stable and precise estimate of *γ*_opt_ with negligible variability across decimals. Connection lengths were defined as the Euclidean distance between the somata of connected neuron pairs, rather than the physical length of neurites. At the whole-brain level, connection lengths were defined as the distance between region centroids. This definition excluded effects of spatially extended axonal or dendritic arbors, isolating instead the probability that two locations separated by a distance *d* form a connection, independent of neurite trajectories.

### Computing whole-brain gradients and eigenmodes

We used a previously published dataset from our lab^6^ to investigate the relationship between whole-brain connectivity gradients and geometric eigenmodes. We used a high-resolution parcellation of 987 nodes per brain hemisphere, obtained by fragmenting larger anatomical brain region masks with k-means clustering, to coarse-grain the activity of roughly 54, 464 ± 5086 neurons per animal (*n* = 22 larvae). Then, we averaged node time series (the first 10 minutes, which correspond to spontaneous activity) from both hemispheres, computed pairwise correlations between nodes, and averaged individual correlation matrices to generate a group-average FC matrix. We calculated FC gradients using the diffusion maps algorithm on the absolute correlations |*C*| with *α* = 0.5 as per the rest of this study. In parallel, we calculated geometric eigenmodes on a 3D whole-brain mask obtained by manually delimiting the external boundaries of the brain. More specifically, we used a cytosolic GCaMP template to properly visualize cell bodies and neuropil regions and segmented the brain boundaries on each 2D plane. Then, we mirrored the segmentations, applied 3D filtering, and binarized again to obtain a smooth 3D mask of the brain outlines. We generated a tetrahedral mesh of this volume and calculated eigenmodes similarly as described previously, then mapped the eigenmode values to the brain region nodes using nearest neighbors before finally correlating both mode ensembles after establishing their optimal mapping.

### Computing brain-wide connectivity

To estimate brain-wide connectivity (Fig.7 **g**), we followed an approach established in prior work^6;43^. Briefly, for each pair of atlas brain regions, defined to be mutually exclusive and collectively exhaustive, we counted neurons whose somas were located in region *i* and whose neurite terminals innervated region *j*. Connectivity weights were obtained by summing these counts across all reconstructed neurons, followed by volume normalization and logarithmic scaling^6^, yielding a weighted, undirected inter-regional projectome. Subsequent analyses were performed on a binarized version of this matrix, retaining only the presence or absence of connections among the 65 brain regions.

### Statistics

Various statistical tests were used throughout the study, depending on the nature of the data and the comparisons being made. In each case, the specific test, sample size, and exact *P* -values are reported in the relevant figure legends or text. All tests were two-sided unless otherwise specified, and significance was defined as *P* < 0.05. No statistical methods were used to predetermine sample sizes. Spatial permutations were performed using the BrainSMASH toolbox^32^, which randomizes data while approximately preserving its spatial autocorrelation as captured by the variogram.

## Data availability

The data used in this study are publicly available in the Université Laval collection on Borealis (https://doi.org/10.5683/SP3/YKCS90). Raw calcium imaging videos are available upon reasonable request to the authors.

## Code availability

Code used in this study is available on https://github.com/DynamicaLab/geometric-networks.

## Acknowledgements

We acknowledge Calcul Québec and Digital Research Alliance of Canada for their technical support and computing infrastructures. We are thankful for discussions with members of the Dynamical Research Lab and the PDK Lab. A.L. and O.R. were supported by PhD scholarships from the Natural Sciences and Engineering Research Council of Canada (NSERC). A.L. was also supported by scholarships from Fonds de recherche du Québec–Nature et technologies (FRQNT), and Unifying Neuroscience and Artificial Intelligence–Québec (UNIQUE). This work was funded by the NSERC (RGPIN-2023-05980 to P.D.K., RGPIN-2024-05626 to A.A., RGPIN-2024-06492 to P.D.), the Sentinelle Nord program of Université Laval (Canada First Research Excellence Fund), the FRQNT–Team Research Project (doi:10.69777/365162 to P.D.K. and P.D.), and a UNIQUE–Neuro-AI Pilot Project grant (P.D.K. and P.D.).

## Author contributions

A.L. and O.R. designed the study. A.L. conducted numerical experiments. O.R. performed mathematical calculations. A.L. conducted calcium imaging experiments and analyzed the data. A.L., O.R. and A.A. contributed to code development. All authors analyzed the data and interpreted the results. A.L. and O.R. wrote the initial manuscript. All authors contributed to manuscript revision. P.D.K., A.A., and P.D. acquired funding and supervised the project.

## Supplementary Material

### Supplementary figures

**Figure S1:**
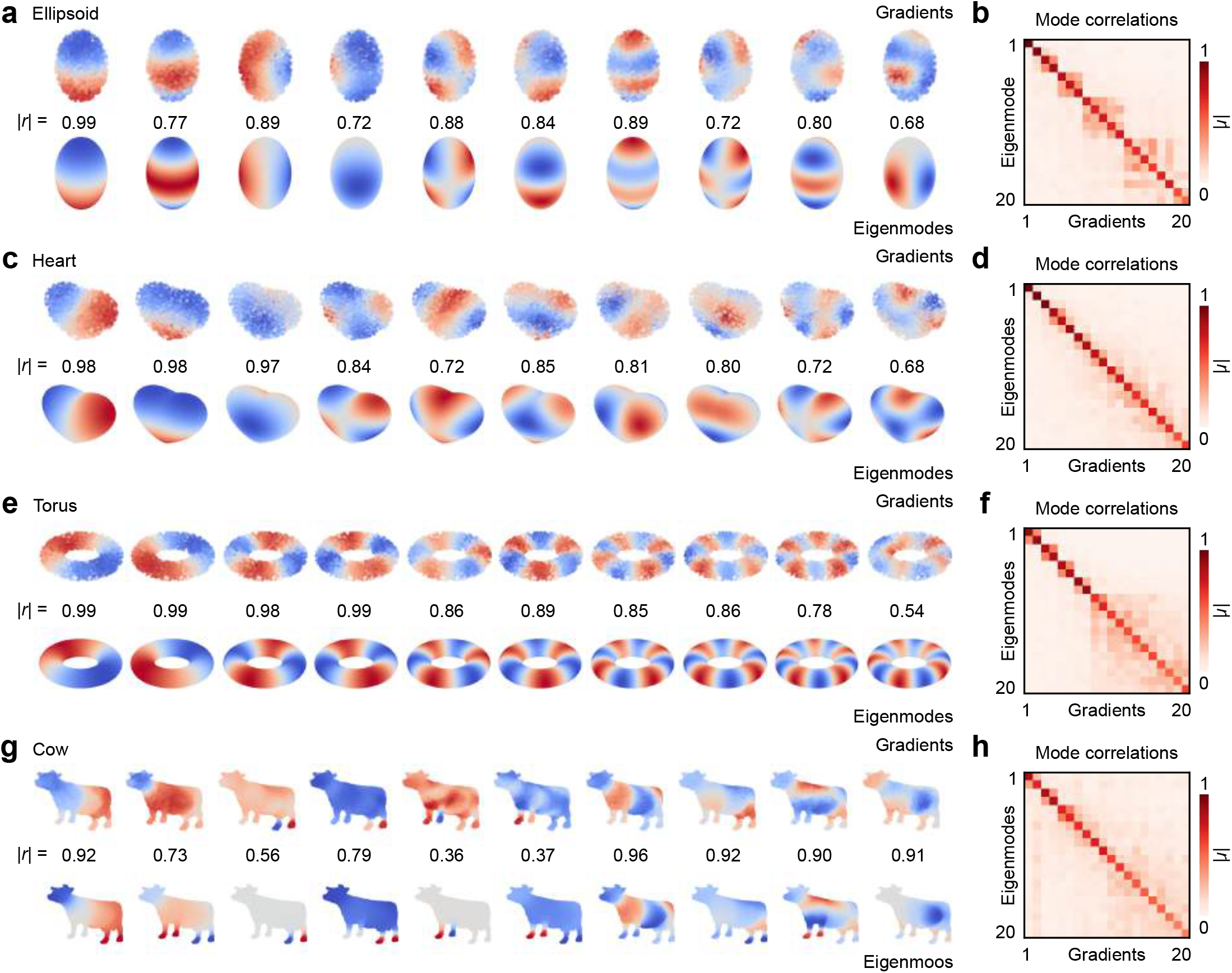
Comparison of functional connectivity gradients and geometric eigenmodes in various geometries. (**a**) Gradients and eigenmodes of an ellipsoid shape; absolute correlations |*r*| are indicated for the first 10 mode pairs (one representative simulation). (**b**) Eigenmode-gradient correlation matrix for the ellipsoid geometry, averaged across 10 coordinate sets. (**c**) Gradients and eigenmodes of a heart shape. (**d**) Eigenmode-gradient correlation matrix for the heart geometry, averaged across 10 coordinate sets. (**e**) Gradients and eigenmodes of a torus shape. (**f**) Eigenmode-gradient correlation matrix for the torus geometry, averaged across 10 coordinate sets. (**g**) Gradients and eigenmodes of a cow shape. (**h**) Eigenmode-gradient correlation matrix for the cow geometry, averaged across 10 coordinate sets. The results shown here were obtained with the EDR model for the first 3 geometries, and the radius model in the fourth geometry due to its better performance in narrower parts of the volume.

**Figure S2:**
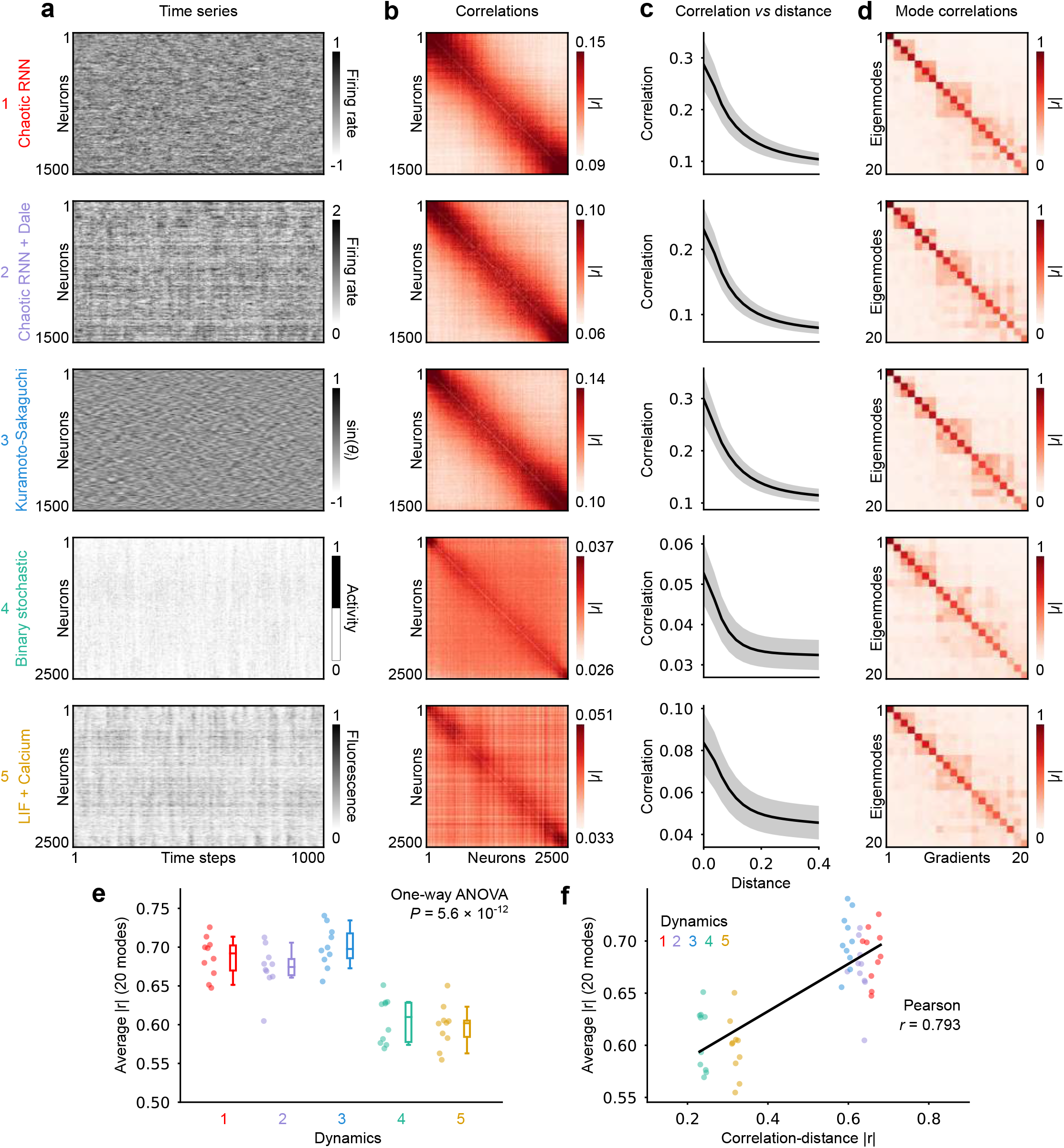
Eigenmode-gradient correlations in ellipsoid for different dynamical systems. (**a**) Example simulated time series from five dynamical models, indicated on the left and detailed in Methods; all simulations are embedded in the ellipsoid geometry with the EDR connectivity rule and *γ* = 27.5. (**b**) Example pairwise neuronal activity correlations for each model, averaged across 50 simulations; the matrix is sorted to follow the *z* axis of the ellipsoid. (**c**) Average correlation-distance relationships for each dynamical model; the shaded region indicates average standard deviation. (**d**) Eigenmode-gradient correlation matrices for each dynamical model, averaged across 10 ellipsoids. (**e**) Average eigenmode-gradient correlations for each dynamical model (10 simulations per model, averaged over mode pairs); different models yield statistically different eigenmode-gradient correlations (one-way ANOVA, *P* = 5.6 *×* 10^−12^). (**f**) Eigenmode-gradient |*r*| as a function of the correlation-distance |*r*| for the five different models; dynamical models whose correlations are more tightly coupled to euclidean distance yield a better alignment with geometry (Pearson *r* = 0.793).

**Figure S3:**
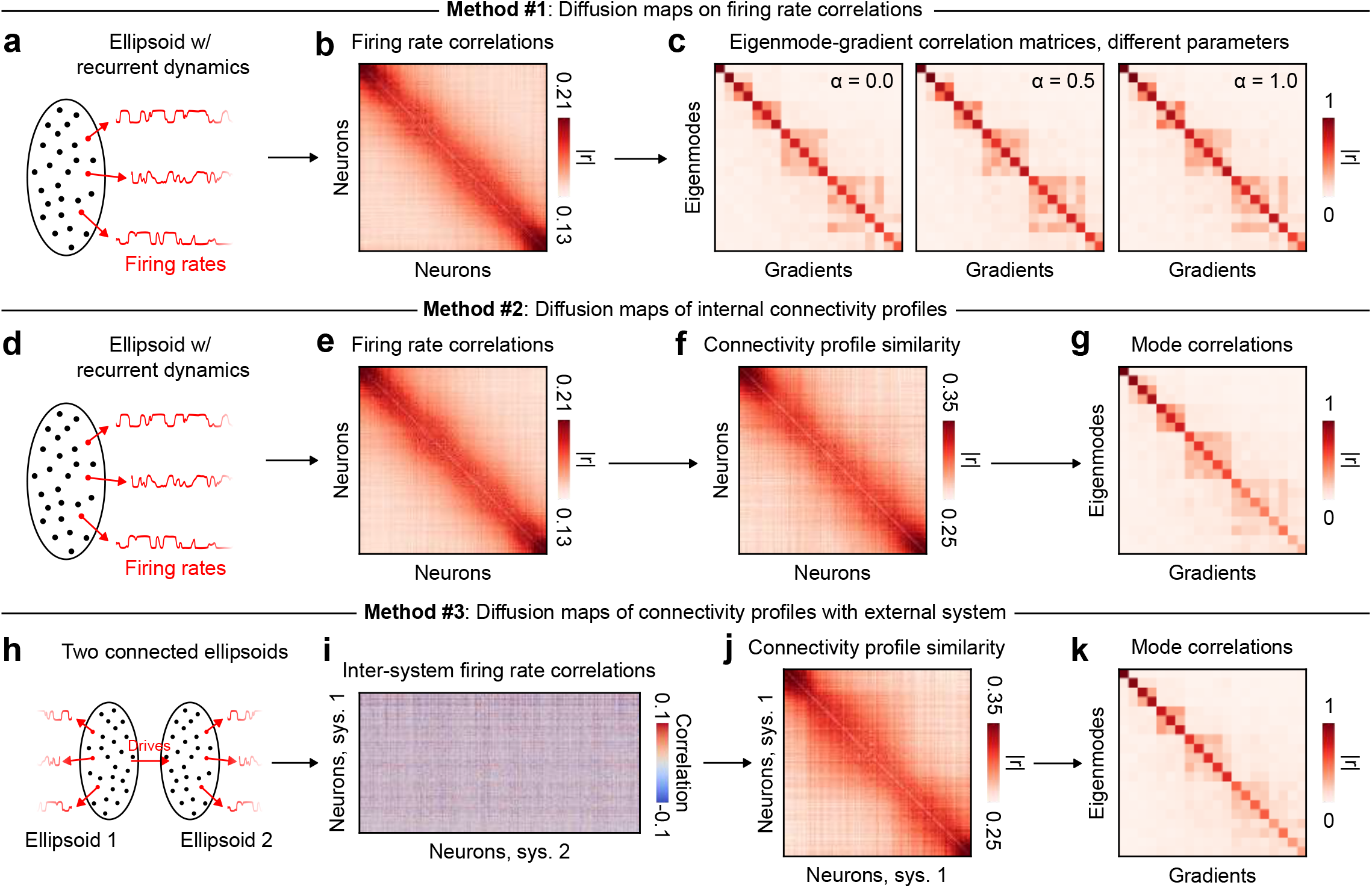
Derivation of geometric gradients using various gradient analysis methods. (**a-c**) Method 1 uses the diffusion maps algorithm on the firing rate correlation matrix to compute gradients, yielding results that are mostly independent of the *α* parameter controlling the effect of local density on the diffusion process. (**d-g**) Method 2 adds one extra step to the first method by computing the connectivity profile similarity from the correlation matrix before computing gradients. (**h-i**) Method 3 approximates the connectopic mapping procedure used in Pang et al^28^, who derived subcortical gradients from their temporal association with cortical activity. Here, two interconnected systems are simulated, one smaller representing a subcortical region, and one larger representing external regions like the cortex. Time series from both ellipsoids are correlated to each other, and then the connectivity profile similarity matrix is used to compute gradients. Overall, this supplementary analysis demonstrates that geometric gradients can be obtained from gradient derivation methods that vary in their sequence of numerical operations, highlighting their methodological robustness.

**Figure S4:**
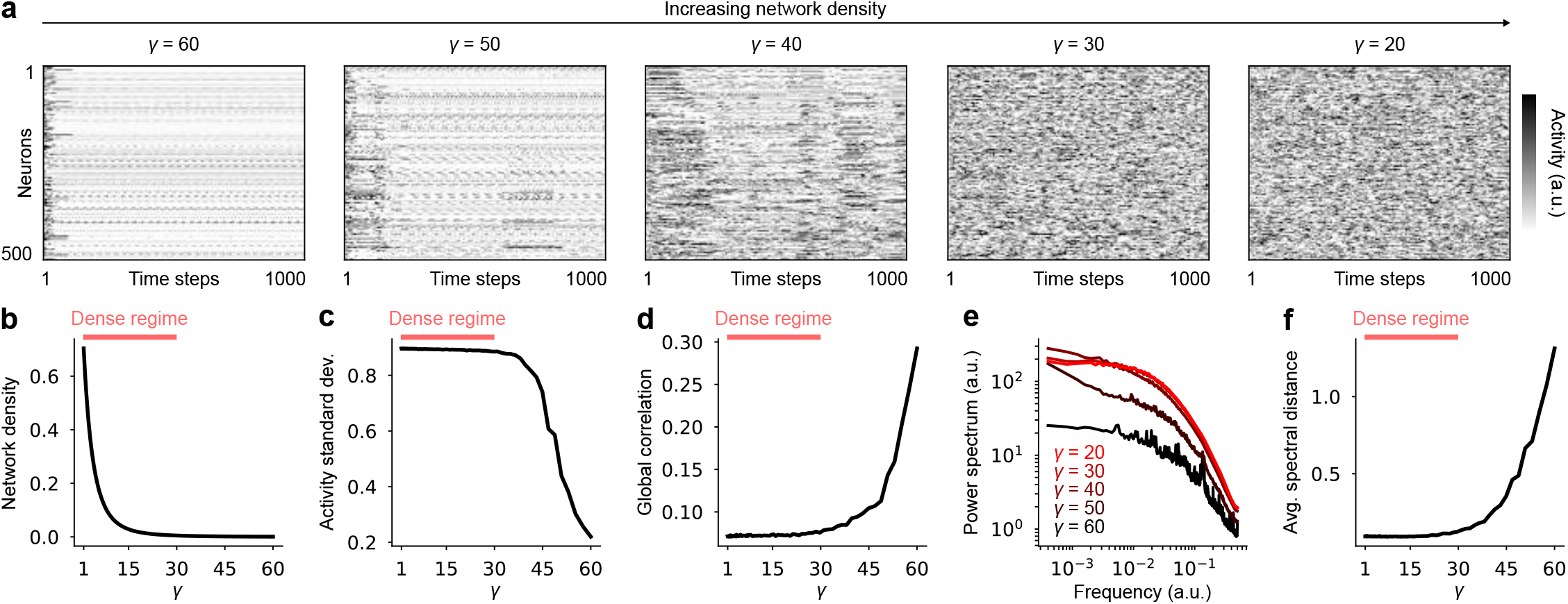
Network dynamics are stable within the dense connectivity regime. (**a**) Example time series from 500 neurons from EDR network simulations at different *γ* values. (**b**) Network density plotted against *γ*. (**c**) Average standard deviation of neuronal firing rates plotted against gamma; standard deviation is small in the sparse regime due to most neurons exhibiting sparse or static activity patterns. (**d**) Average absolute correlation between neurons plotted against *γ*; global correlations increase in the sparse regime due to periodic oscillations or steady-state dynamics. (**e**) Average power spectra at 5 different *γ* values, indicated on the plot; dynamics on denser graphs converge toward power-law spectra, whereas sparse graphs yield peaky spectra indicating the presence of more periodic oscillations. (**f**) Average spectral distance plotted against *γ*; spectral distance is measured between all pairs of power spectra measured at different *γ* values, then averaged, thus indicating how one spectrum resembles all other spectra; lower distances in the dense regime suggest that spectra have converged toward stable power laws. In (**b, c, d, f**), black curves are obtained by averaging over 10 simulations of *N* = 2500 neurons per *γ* value within the ellipsoid geometry. A pink rectangle roughly highlights the “dense” regime (*γ* < 30), in which most analyses of the paper are conducted. Dev., deviation; Avg., average.

**Figure S5:**
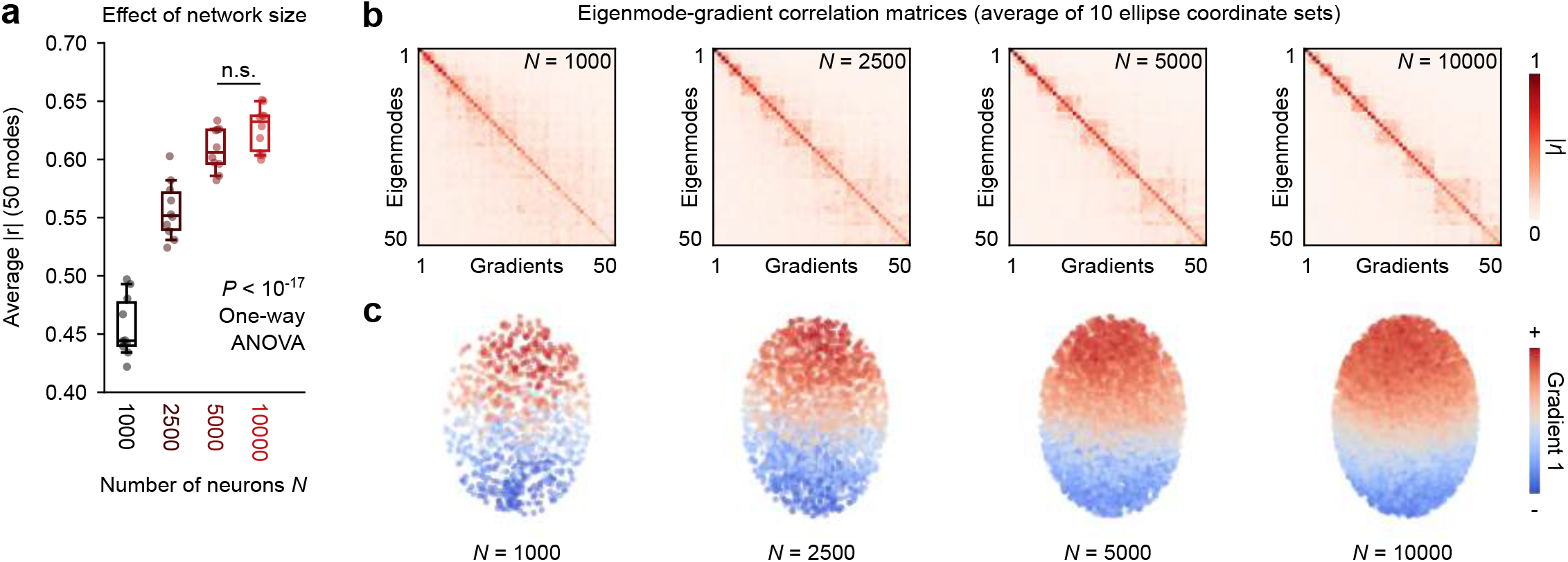
Eigenmode-gradient correlations in ellipsoid for different network sizes. (**a**) Average eigenmode-gradient correlation (averaged over 50 mode pairs) for four different network sizes; denser networks exhibit gradients that correlate more with geometric eigenmodes (one-way ANOVA, *P* < 10^−17^). (**b**) Eigenmode-gradient correlation matrices for different network sizes (indicated in the top right matrix corners), averaged over 10 ellipsoids. (**c**) Neuron centroids colored by the numerical values of the first functional gradient for increasing network sizes.

**Figure S6:**
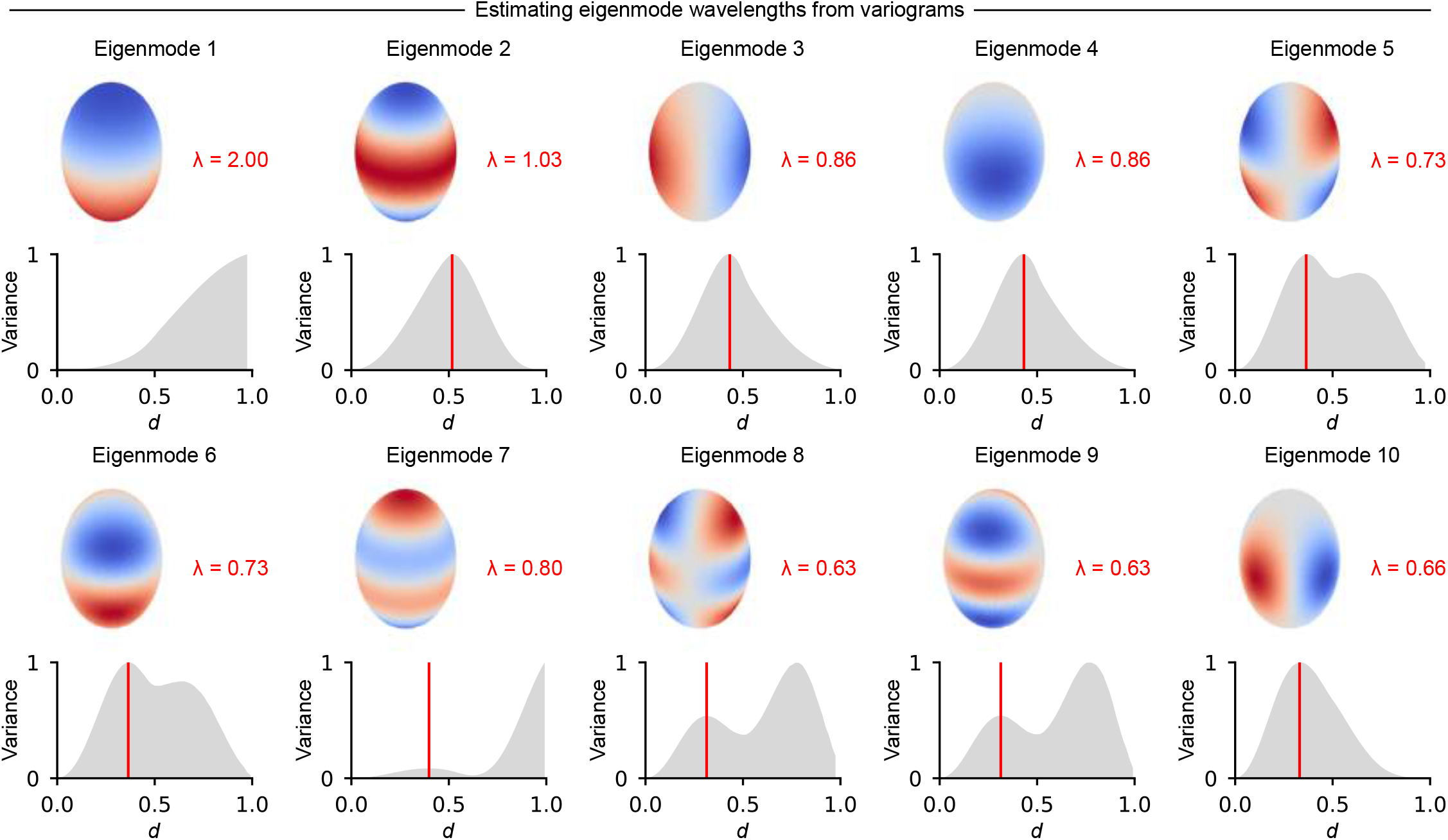
Estimating eigenmode wavelengths using their spatial variograms. The first 10 ellipsoid eigenmodes are plotted alongside their wavelengths (top rows), which are estimated from the first peak of each variogram (bottom rows, Methods).

**Figure S7:**
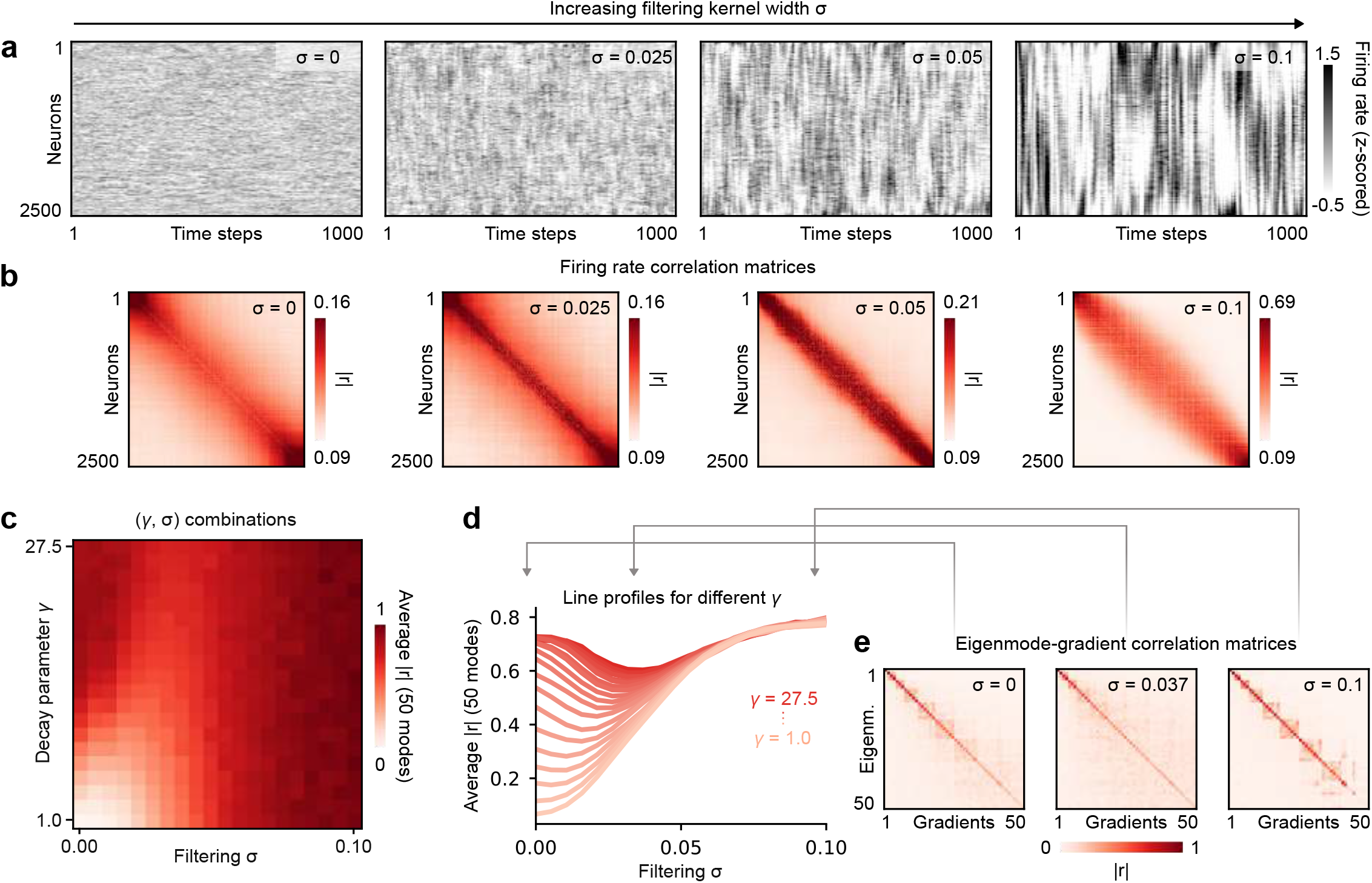
Combined effects of local connectivity and spatial filtering in numerical simulations. (**a**) Examples of simulated firing rate time series (*N* = 2500 neurons, EDR connectivity) for increasing levels of spatial filtering (from left to right); neurons are sorted along the major axis of the ellipsoid to accentuate patterns. (**b**) Pairwise correlations of neuronal firing rates at increasing levels of spatial filtering, associated with the four time series matrices of **a**; colors are scaled to the 5th and 95th percentiles of correlations to improve visualization. (**c**) Grid of eigenmode-gradient correlations averaged over 50 mode pairs for different values of *γ* and *σ*; *N* = 2500 neurons, 10 ellipsoid coordinate sets per pixel, with 50 averaged simulations per set. (**d**) Rows of the matrix of **c** with mild gaussian filtering to improve readability. (**e**) Eigenmode-gradient correlation matrices for *γ* = 27.5 and three different values of *σ*; the middle matrix corresponds to the case where filtering weakened the eigenmode-gradient correlations. Interestingly, spatial filtering has a non-monotonous effect in the most locally connected networks, initially decreasing the alignment with geometry before increasing again.

**Figure S8:**
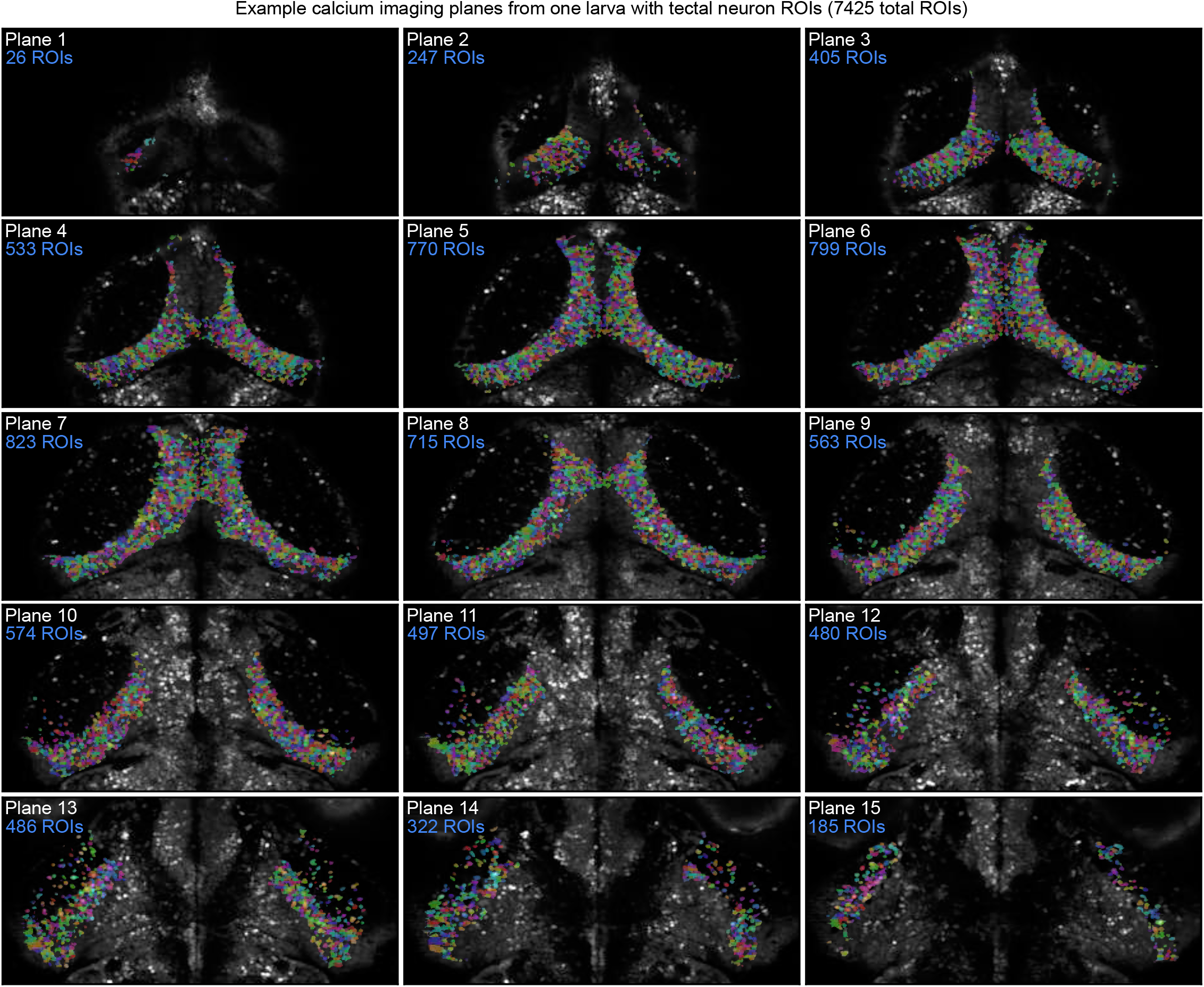
Temporal averages of two-photon calcium imaging planes from one example animal, with pseudocolored nuclei located within the optic tectum identified using Cellpose 2.0. The number of detected regions of interest (ROIs) is indicated on each plane. Imaging volumes extend from the most dorsal (top left) to the most ventral (bottom right) parts of the tectum across both hemispheres to ensure that the entire 3D structure is properly sampled.

**Figure S9:**
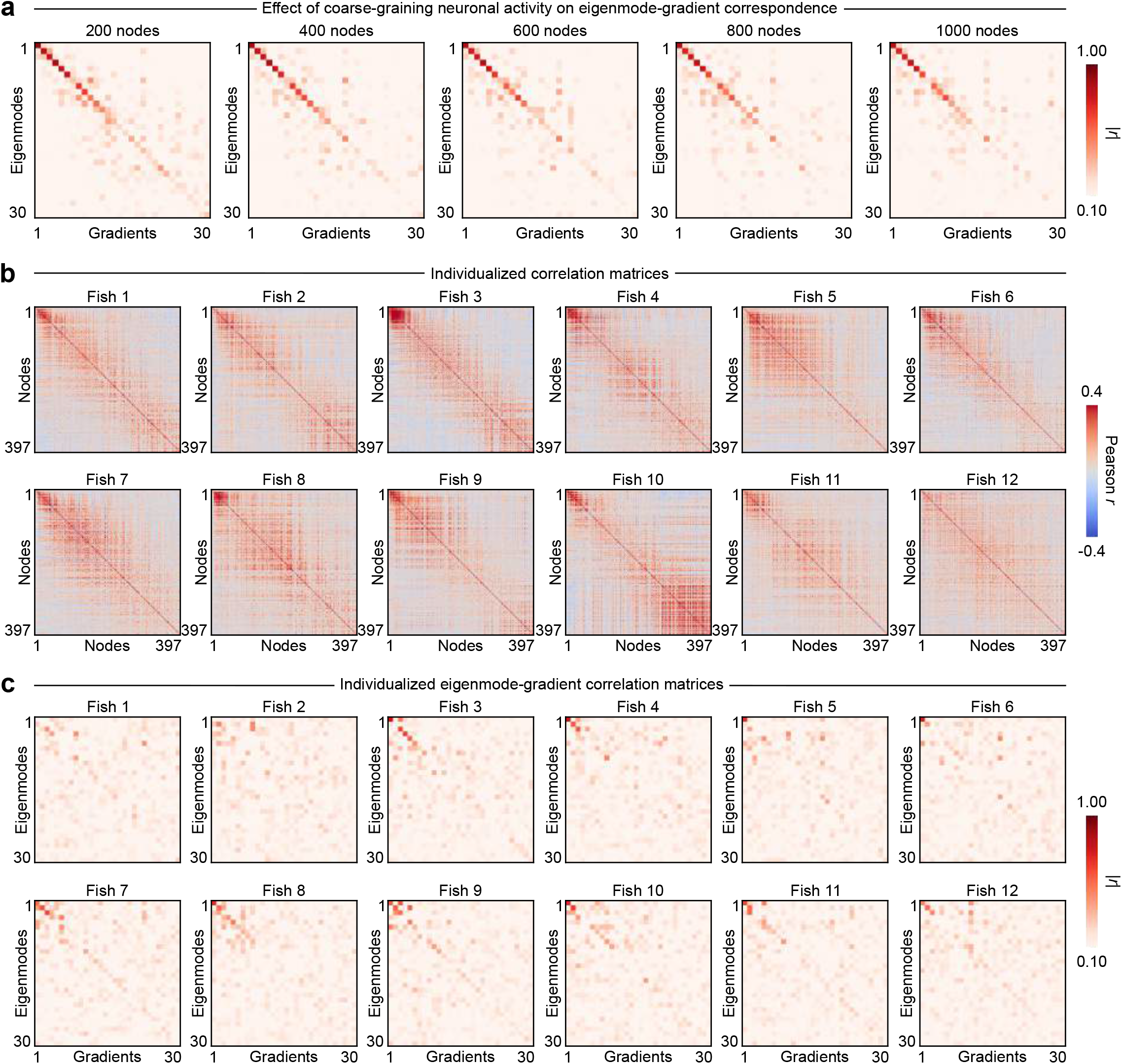
Effect of coarse-graining and averaging on geometric gradients. (**a**) Eigenmode-gradient correlation matrices for varying numbers of network nodes in the optic tectum; notice the minimal effect of spatial resolution on the eigenmode-gradient correlations. (**b**) Correlation matrices of optic tectum node activity for 12 different larvae, sorted from anterior to posterior nodes. (**c**) Eigenmode-gradient correlation matrices calculated on the individual correlation matrices from the previous panel; notice the poor quality of the correspondence in most animals. This suggests that individual imaging sessions insufficiently sample the correlation structure of the optic tectum.

**Figure S10:**
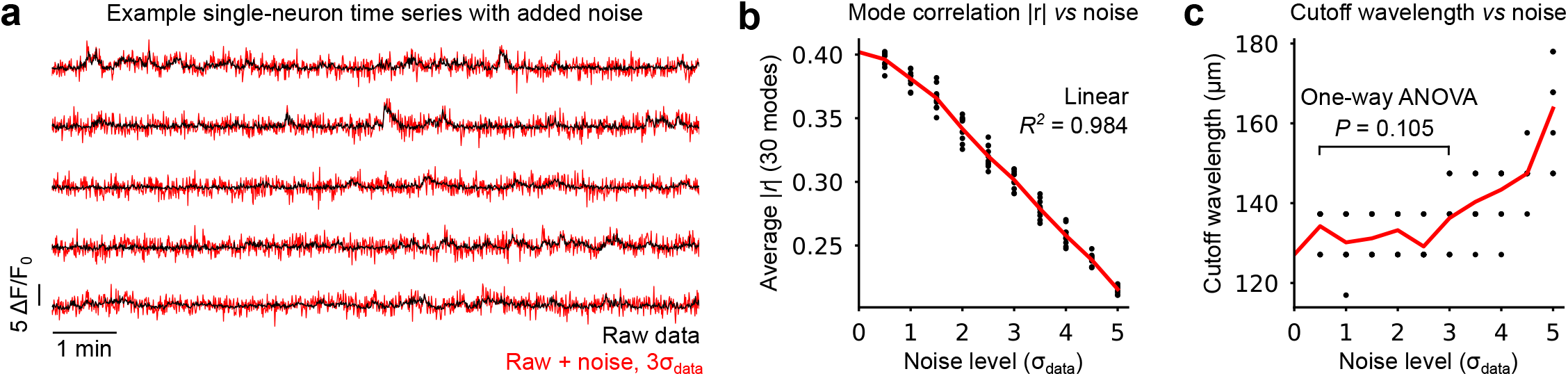
Influence of experimental noise on the geometric cutoff wavelength. (**a**) Five example single-neuron fluorescence time series, with varying noise levels; black, raw traces; red, raw traces with numerical noise drawn from a normal distribution with 3 times the standard deviation of the data. (**b**) Absolute correlation |*r*| averaged across 30 eigenmode-gradient pairs for varying noise levels applied to fluorescence time series; red, average curve; black dots, 10 replicates per noise level. (**c**) Cutoff wavelength inferred from a piecewise linear fit on eigenmode-gradient correlation matrix diagonals for varying noise levels (see Methods); red, average curve; black dots, 10 replicates per noise level; despite a gradual decrease of correlations observed in **b**, no significant cutoff wavelength differences are observed until noise levels exceed 3*σ*_data_ (One-way ANOVA, *P* = 0.105, *F* = 1.929). This analysis suggests that reasonable levels of experimental noise have a negligible influence on the position of the cutoff point in the eigenmode-gradient mapping.

**Figure S11:**
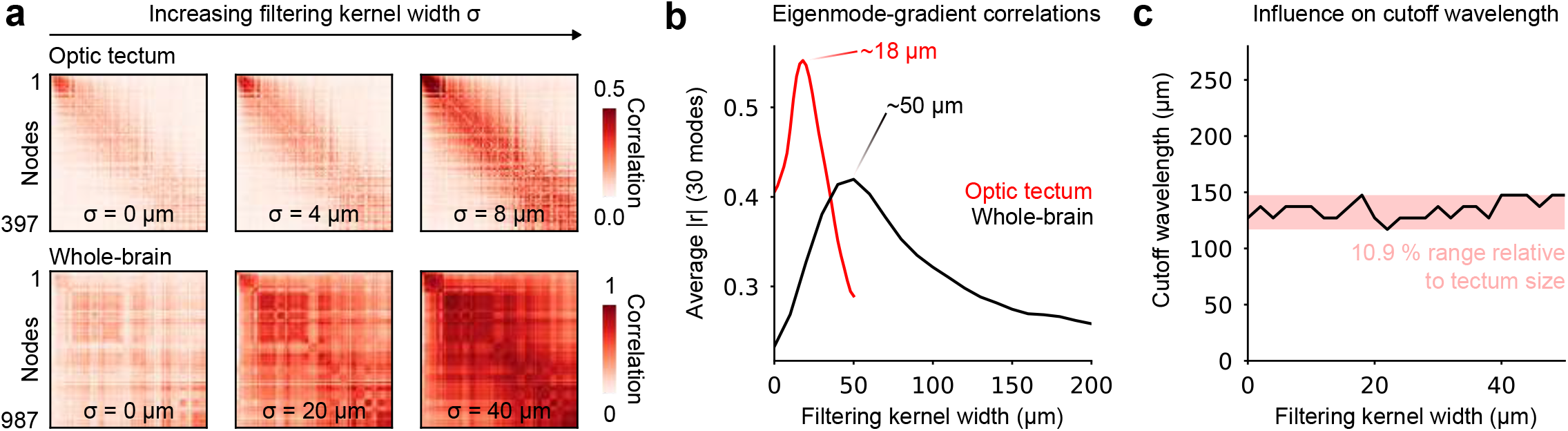
Spatial filtering drives geometric eigenmodes in both optic tectum and whole-brain calcium imaging data. (**a**) Tectal (top) and whole-brain (bottom) FC matrices for varying spatial filtering kernel sizes, indicated on each matrix (*n* = 12 larvae for tectal data, *n* = 22 larvae for whole-brain data). (**b**) Eigenmode-gradient correlation |*r*| averaged over 30 mode pairs for the optic tectum (red) and the entire brain (black) at varying filtering kernel sizes; optima are indicated for each curve. (**c**) Cutoff wavelength inferred using a piecewise linear fit on the mode correlation matrix diagonal at varying filtering levels (optic tectum only, see Methods). Spatial filtering does not change the cutoff wavelength beyond a ∼ 10% margin.

### Supplementary Mathematical Note

#### Calculation of filtering-induced correlations

We demonstrate the claim made in the main text that the expected correlations induced by filtering uncorrelated gaussian noise signals are given by

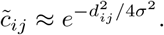

Let us begin with the Pearson correlation coefficient

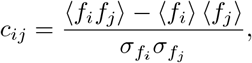

where *f*_*i*_ is the random variable corresponding to the signal measured from node *i*, ⟨*f*_*i*_⟩ is its expected value, and 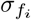 is its standard deviation. Without loss of generality, we assume that ⟨*f*_*i*_⟩ = 0 for all *i*, which implies that

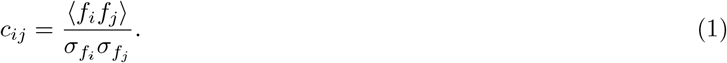

The spatial filtering considered in the main text consists in the convolution of the activities with a Gaussian filter, meaning that the signal measured from node *i* becomes

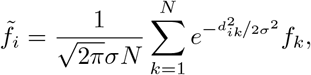

where *σ* is the standard deviation of the Gaussian filter, *N* is the number of nodes in the network, and *d*_*ik*_ is the distance between nodes *i* and *k*. The covariance between these filtered signals is

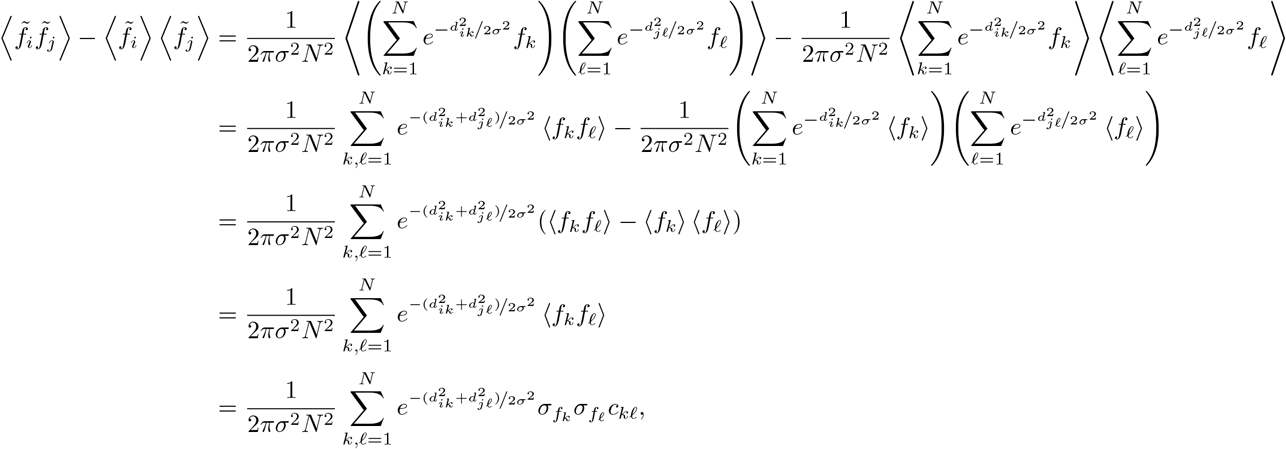

where we used the fact that ⟨*f*_*i*_⟩ = 0 for all *i*, and invoked Eq. (1). The variance of each filtered signal simplifies to

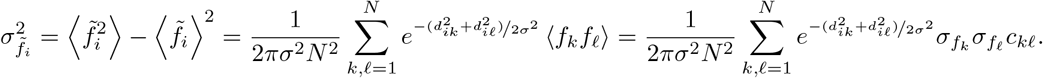

The correlation between the filtered signals therefore is

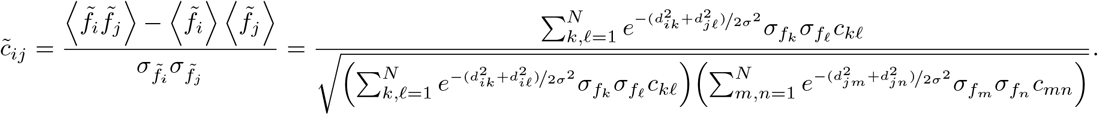

The denominator is superfluous, since 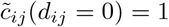 by definition, meaning it can simply be obtained through normalization.

#### Independently and identically distributed signals

We now assume that the raw signals are independently and identically distributed, which further implies that 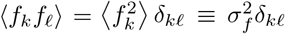 and that *c*_*kℓ*_ = *δ*_*kℓ*_, where *δ*_*kℓ*_ is the Kroenecker delta. The correlation between the filtered signals simplifies to

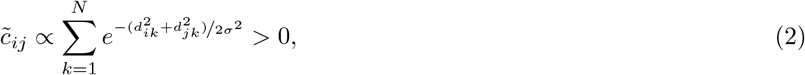

which demonstrates that filtering uncorrelated signals does induce correlations.

Exact computation of 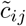 depends on the distribution of the distances between the nodes and on the precise geometry of the manifold, and cannot be carried out analytically further in general. However, assuming that nodes are distributed uniformly in the manifold, and ignoring the effect of the manifold’s boundaries, we can use a continuous approximation which becomes exact as *N* → ∞.

Under these assumptions, combined with the law of cosines

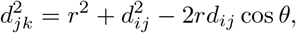

and with the following geometric construction,

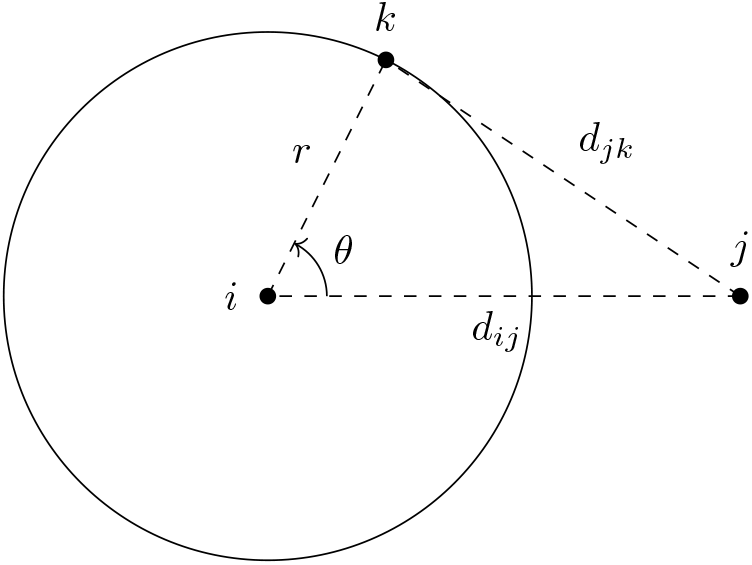

the numerator in Eq. (2) can be approximated as

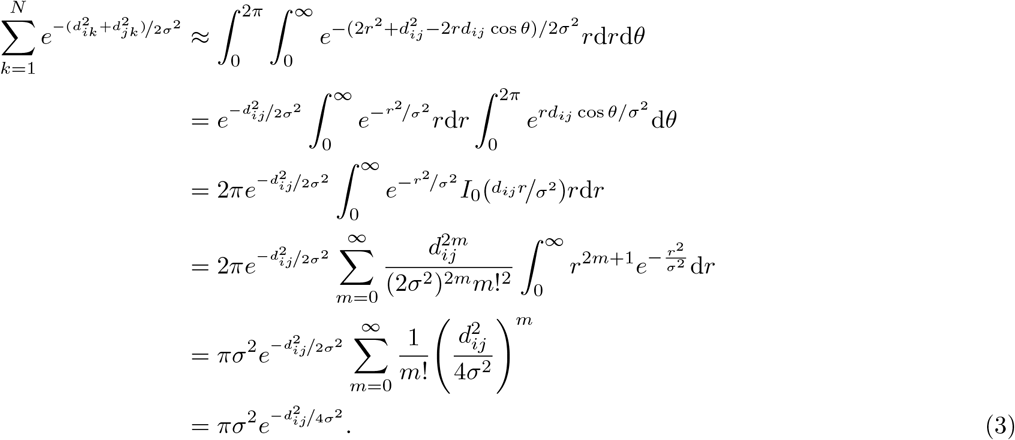

Normalizing at *d*_*ij*_ = 0 yields

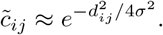

#### Correlated and identically distributed signals

In the case where the signals are identically distributed, but are correlated, the general formula

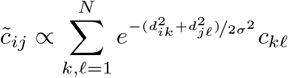

does not lend itself to further simplification, unless the correlations follow a Gaussian profile

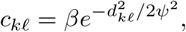

in which case

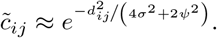

Note that we correctly retrieve the independent case for *ψ* = 0, as well as the original correlations, *c*_*kℓ*_, up to constant factor *β* when the width of the filtering kernel vanishes (*σ* = 0), as expected.

Finally, gaussian smoothing of correlations following an exponential profile with parameter *γ* yields

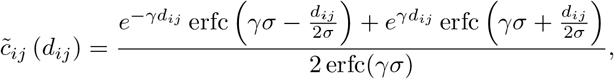

which is simply the convolution of the exponential with two gaussians of variance *σ*^2^.

